# Data-guided Multi-Map variables for ensemble refinement of molecular movies

**DOI:** 10.1101/2020.07.23.217794

**Authors:** John W. Vant, Daipayan Sarkar, Ellen Streitwieser, Giacomo Fiorin, Robert Skeel, Josh V. Vermaas, Abhishek Singharoy

## Abstract

Driving molecular dynamics simulations with data-guided collective variables offer a promising strategy to recover thermodynamic information from structure-centric experiments. Here, the 3-dimensional electron density of a protein, as it would be determined by cryo-EM or X-ray crystallography, is used to achieve simultaneously free-energy costs of conformational transitions and refined atomic structures. Unlike previous density-driven molecular dynamics methodologies that determine only the best map-model fits, our work uses the recently developed *Multi-Map* methodology to monitor concerted movements within equilibrium, non-equilibrium, and enhanced sampling simulations. Construction of all-atom ensembles along chosen values of the Multi-Map variable enables simultaneous estimation of average properties, as well as real-space refinement of the structures contributing to such averages. Using three proteins of increasing size, we demonstrate that biased simulation along reaction coordinates derived from electron densities can serve to induce conformational transitions between known intermediates. The simulated pathways appear reversible, with minimal hysteresis and require only low-resolution density information to guide the transition. The induced transitions also produce estimates for free energy differences that can be directly compared to experimental observables and population distributions. The refined model quality is superior compared to those found in the Protein DataBank. We find that the best quantitative agreement with experimental free-energy differences is obtained using medium resolution (~5 Å) density information coupled to comparatively large structural transitions. Practical considerations for generating transitions with multiple intermediate atomic density distributions are also discussed.

## I. INTRODUCTION

Single-particle cryo-electron microscopy (cryo-EM) has evolved into one of the most effective structure determination tools in modern-day structural biology. Following advances in electron detector technology,^1^ cold field-emission electron gun sources and energy filters,^2^ cryo-EM has achieved resolutions rivaling those of X-ray crystallography or NMR spectroscopy,^3^ often providing novel structures or conformations.^4–6^ However, static X-ray or cryo-EM structures alone offer limited information on the function of biomolecules. The determination of conformational trajectories remains a key stumbling block towards associating structure and function. These trajectories are expected to deliver substantial information beyond static structures, revealing, for example, the propagation of allosteric signals in complex biological molecules,^7^ and important clues to the conformational diversity of sites associated with diseases.^8,9^

Traditionally, molecular trajectories are derived using molecular dynamics (MD) simulations imposing either classical assumptions via all-atom and coarse-grained force fields,^10,11^ or by introducing *ab initio* methodologies coupled with the classical particles via extended Lagrangian schemes.^12–14^ However, it is now well established that biologically relevant conformational transitions and timescales remain inaccessible to brute force MD. This drawback of traditional MD has motivated the inception and application of a range of alchemical^15,16^ and geometric methods^17–23^ for enhanced sampling of molecular movements.

Experimental methods have also moved beyond calorimetric measures to capture the thermodynamic manifestations of structural ensembles and molecular trajectories. Single-molecule measurements routinely derive free energy profiles and rates as a function of simple distance or angular metrics,^24,25^ though their spatial resolution is limited. Highly-resolved molecular ensembles are determined from NMR, and EPR experiments,^26,27^ but such data is limited in size compared to MD and typically misses fast kinetic information. Addressing the need to construct free energy surfaces directly from experiments while simultaneously recovering the conformational changes, geometric machine learning methodologies are employed to hierarchically cluster millions of 2dimensional single-particle images onto a low-dimensional manifold using diffusion maps.^28^ The population of points on this manifold is correlated to free energy changes between 10 to 100 molecular conformations by a Boltzmann factor.^29^ Such examination of the conformational trajectories (the so-called “molecular movies”) from the cryo-EM data offers arguably the first experimentally-verifiable and structurally-resolved view of an entire free energy landscape, including both the intermediates and rare conformations. Therefore, going beyond the visualization of realistic stationary structures, incorporating these energy-ranked cryo-EM ensembles in MD can accelerate the potential of mean force (PMF) estimation from simulations.

Integration of cryo-EM data, and more generally, experimental data with MD, has followed from the development of two families of methods, namely flexible fitting,^30,31^ and Bayesian inferencing.^32^ While the former serves as a real space refinement tool available in almost all the structure determination software,^33,34^ the latter has been successful in either folding small proteins (< 115 residues)^32^ or seeking small-scale structural changes of subdomains (<5 Å of root mean square deviation (RMSD)) and free energy changes within larger cryo-EM density segments.^35^ The combination of flexible fitting and Bayesian inferencing^36^ overcomes this system-size restriction on protein folding and captures extremely large-scale conformational transitions from cryo-EM data of heterogeneous complexes. Nevertheless, extracting the free energy from these integrative simulations is non-trivial. A reduced representation based on collective variables would lend itself to the computation of PMFs to be compared with experiments.

As a step towards facilitating data-guided free energy estimations, we propose here to use directly the 3D electron density fields to define plausible reaction coordinates. To this end, we employ the recently introduced *Multi-Map* method,^37^ which uses volumetric maps to measure and simulate changes in shape for molecular aggregates. This is achieved by quantifying the similarity between the instantaneous molecular configuration and each of the target volumetric maps. The Multi-Map method has so far been successful at computing the thermodynamic cost of wetting/dewetting in hydrophobic cavities,^37^ as well as membrane deformations that are either spontaneous^37^ and protein-induced.^38^ It is thus tempting to use this method to simulate changes in the internal structure of biological macromolecules.

Here, we demonstrate that the use of volumetric maps representing cryo-EM densities resolved between 1-9 Å, allows modeling of large-scale transformations in protein structure. Biased sampling along Multi-Map variables constructed from this density induces the protein to alter its shape reversibly in the manner prescribed by the series of electron densities, and the corresponding PMF is derived from the simulated trajectory. Typically, high-quality atomistic structures for each of the relevant states are required to simulate a molecule’s transformation and extract the associated PMF. However, formulating the free energy problem in terms of the density itself enables the attainment of structural ensembles by biasing any starting model to a given state defined by a density map. By varying the maps’ resolution between the atomic and molecular scale, we allow simultaneous real-space refinement of the data and biased sampling of the conformations. Thus, starting with only one high-quality atomistic structure defining one end state in a series of maps, low-resolution EM maps corresponding to the adjacent states can be sampled producing refined atomistic models for all the states. Due to such built-in refinement capability, free energy surfaces can be obtained even from Multi-Map variables based on low-resolution maps, with accuracy consistent with those from high-resolution structures and maps.

In what follows, the conformational dynamics of three protein molecules are investigated: apo and AP_5_A-bound adenylate kinase (ADK), carbon monoxide dehydrogenase (CODH) and *Francisella* lipoprotein3 (FLPP3). To allow comparisons between the three, synthetic density maps of equal resolutions were generated, and used to construct a Multi-Map collective variable for each protein (Sect. II.B). The proteins are then simulated with equilibrium MD (Sect. III.A), nonequilibrium MD (Sect. III.C) and enhanced sampling simulations (Sect. III.D).

An analysis of the non-equilibrium work associated with these conformations offers a theoretical framework to determine the “resolvability” of a map.^39^ Employing two maps for each protein, we demonstrate that a two-state Multi-Map variable is able to monitor open and closed protein conformations in equilibrium and during slow conformational transitions, and how the accuracy of the free energy estimates changes with the density map resolution. Finally, the effect of solvation environments on the PMF is discussed, and limitations in capturing nominal structural changes, such as single sidechain rearrangements, are brought to light.

## II. METHODS

To allow quantitative comparison between the three proteins studied and explore the role of density map resolution, we generated synthetic maps from atomic models with multiple states deposited in the Protein Data Bank (PDB). These maps were then used to construct a Multi-Map variable^37^ for enhanced sampling. The conformational dynamics of A→B and the reverse B→A transition were monitored using the Multi-Map variable itself, the Euclidean distance between states as measured by root mean square deviation (RMSD), sidechain contacts and crosscorrelation (CC) to the target densities. In addition to varying the system-sizes and environmental conditions, we compared the results obtained from maps generated at five different resolutions (1, 3, 5, 7, and 9Å).

### A. Molecular Dynamics Simulations

Modeling and system setup utilized VMD,^40^ leveraging the *solvate* and *autoionize* plugins to add a 20 Å water layer and a neutralizing concentration of NaCl. All simulations described below share standard simulation parameters consistent with the CHARMM36m and TIP3P force fields used to describe the protein, ions and water.^41,42^

Explicit solvent simulations were run with 2 fs timesteps enabled by restraining bond lengths to hydrogen atoms via the SETTLE algorithm,^43^ and a 12Å cutoff, which is switched at 10 Å. Temperature for all simulations was maintained by a Langevin thermostat set to 300 K, and pressure was maintained by a Langevin barostat set to 1 atm.^44,45^ Long-range electrostatics were calculated using particle mesh Ewald with a 1 Å grid spacing.^46^ The parameters for vacuum, Generalized Born Implicit Solvent (GBIS), and explicit solvent simulations are outlined in Table S1. Equilibrium MD simulation parameters for these systems were carried out with the molecular dynamics engine NAMD 2.14b1.^47^

### B. Construction of the Multi-Map Collective Variable

The recently developed Multi-Map variable^37^ is briefly summarized here. Given the Cartesian coordinates of *N* atoms of interest, indicated as **R** = **r**_1_, **r**_2_,…,**r**_*N*_ with **r** = (*x,y,z*), and *ϕ_k_* (*k* = 1…*K*) a set of volumetric maps, the general form of a Multi-Map variable *ζ* is:

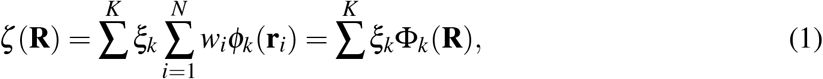

where *w_i_* is the statistical weight assigned to the *i*^th^ atom, and *ξ_k_* represents the contribution of the atomic configuration **R** to the state **Φ**_*k*_ along a *K* points-long pathway.^37^ The physical nature of this pathway is thus determined, aside from the assigned statistical weights, by the choice of the maps themselves.

In the following, we assume each map *ϕ_k_* (**r**) to represent the electron density of a protein, as it would be determined from a cryo-EM or crystallography experiment. For each 3D electron density, the value of the corresponding map *ϕ_k_*(**r**) varies between a maximum value *ϕ_max_* and a threshold value *ϕ_thr_*, as done in the traditional implementation of the MDFF method.^48^ The use of *ϕ_thr_* is dictated by the use of an experimentally measured map, and the choice of its value is simplified by the analysis of the cryo-EM density histogram with the MDFF plugin in VMD.^40,49,50^ Generally, cryo-EM maps will display a large density peak corresponding to the solvent; a threshold value at or above the solvent peak should be chosen to yield a flat potential in the solvent regions (see Wells et al. for a detailed discussion).^51^

To confine sampling to the transition between two states defined by cryo-EM maps, we start with two maps (*ϕ_A_* and *ϕ_B_*), and we choose the coefficients *ξ_A_* = −1 and *ξ_B_* = +1. With this choice, the Multi-Map colvar in Eq. 1 is a two-state variable:

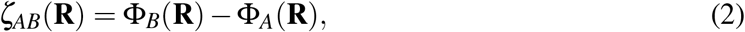

where **Φ**(**R**) = ∑_*i*_*w_i_ϕ*(**r**_*i*_) measures the fitness of the atomic configuration **R** against the threedimensional map *ϕ*(**r**). The two-state colvar *ζ_AB_* lies on a range between a minimum negative and maximum positive value, which correspond to perfect fits to maps A and B, respectively. The range of *ζ_AB_* be estimated *a priori* using the formulation described in Appendix A. Since this range can vary depending on the specifics of the system and the map resolution, during analysis we frequently re-normalize this range into a reaction progress coordinate (changing between −1 and +1) from states A→B.

Fig. 1 illustrates how the Multi-Map variable *ζ* is constructed to link density maps of multiple protein states to conformational transitions, with the specific setup provided in Appendix B. The use of a two-state variable *ζ_AB_* thus defined also draws upon the formalism of other two-state paradigms, such as two-state RMSDs and anisotropic networks,^54,55^ where the two endpoints are known, and the transition between them is simultaneously monitored from both the pathway termini. In analogy, when *ζ_AB_* is at its minimum, **R** is fitted to cryo-EM map A; while when *ζ_AB_* is at its maximum the structure, **R**, is fitted to cryo-EM map B (Fig. 1). *ζ_AB_* values near 0 represent protein configurations neither in state A or in state B. The conformational space near 0 is vast, necessitating a thorough sampling of the associated cartesian space to determine any statistical average. However, the advantage of the two-state *ζ_AB_* sampling protocol is that its gradient is steepest along the most direct path between state A and B. The number of these reactive conformations is much smaller than those needed to be monitored in a protocol using as variable either Φ_*A*_-only or Φ_*B*_-only, where a productive exit from A does not guarantee entry in B, and therefore the system would need to sample an intractably larger phase space to obtain a free energy estimate.

**Figure 1.**
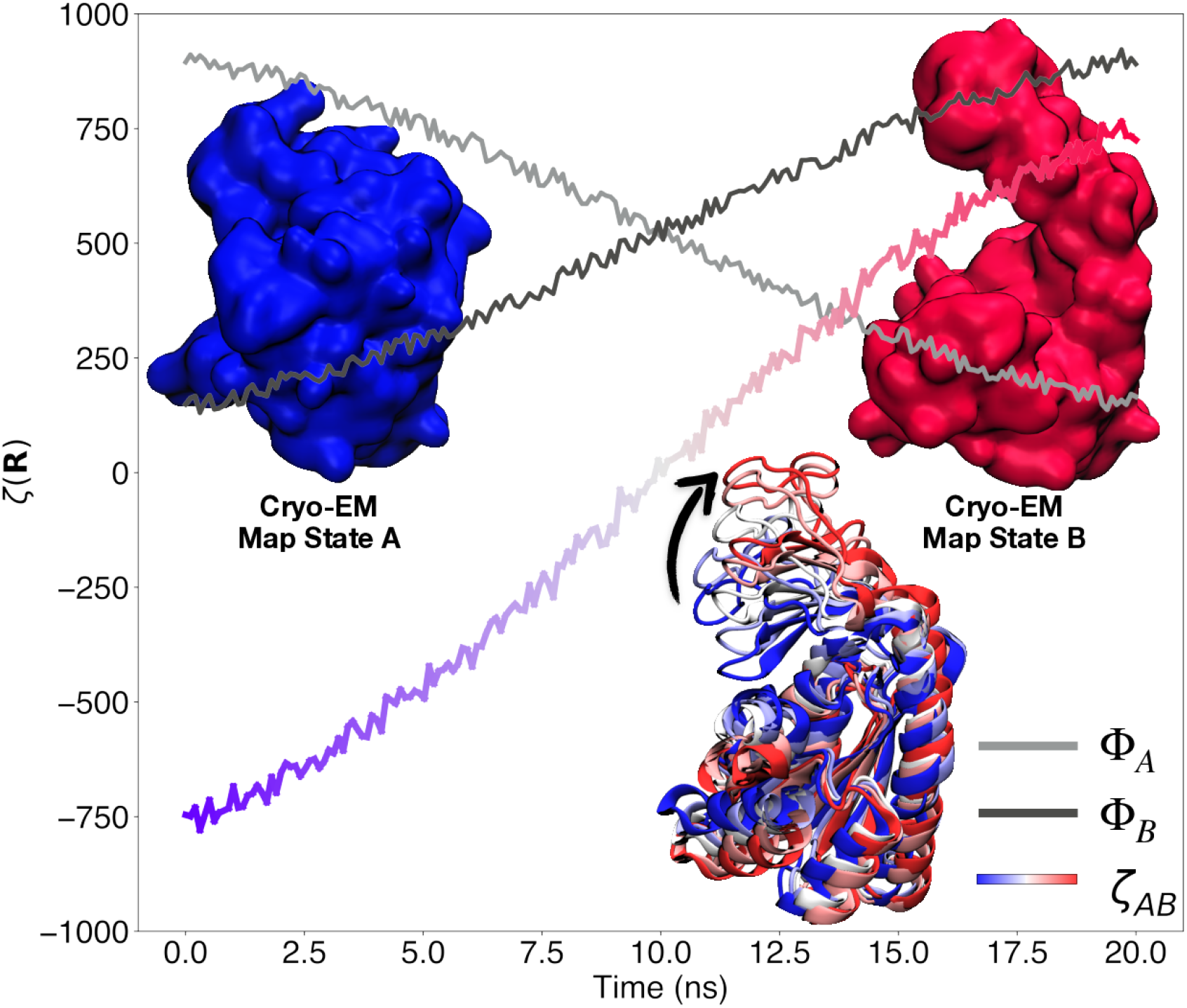
Illustation of three possible choices of Multi-Map reaction coordinates capturing the closed to open transition of ADK. Where Φ_*A*_ and Φ_*B*_ describes a structures similarity to cryo-EM maps state A and B, respectively (*K* = [A] and *ξ_A_* = 1 in Eq. 1 for state A and *K* = [B] and *ξ_B_* = 1 for state B). The variable *ζ_AB_* (Eq. 2) has a negative value for structures similar to state A and a positive value for those like state B. The bottom of the plot shows snapshots of ADK transitioning from state A to B color-coded by the value of the *ζ_AB_* collective variable. The arrow indicates the direction of motion the LID domain of ADK takes during the transition. Surface representations for cryo-EM maps corresponding to states A and B of ADK (PDB IDs 1AKE^52^ and 4AKE,^53^ respectively).

Generalization of the two-state colvar for incorporating more than two cryo-EM maps could benefit from existing computational methodologies to analyze density maps. If the sequence of events captured by the *K* maps is predetermined e.g., by machine learning,^56^ the overall conformational transition can be captured simply by concatenating the 2-state transformations along the pathway.^57^ If the sequence of events is unknown, then combinations of these two state transitions will have to be repeated following different orders of the events until the lowest energy or work pathway is determined for subsequent refinement.^58^

### C. System Preparation

Synthetic density maps were constructed for the demonstration of the Multi-Map colvar. The structures representing states A and B for each system are shown in Fig. 2, and were the density targets used to drive transitions between states. First, molecular systems were chosen based on having multiple conformational states for a single structure in the Protein Data Bank.^59^ The proteins used in this study and their corresponding PDBIDs are shown in Fig. 2. These systems were translated into simulatable models through psfgen, using the CHARMM36m protein force field^41^ and the compatible TIP3P water model.^42^ Second, a map corresponding to a specific state, was generated using the `mdff sim` command, which is part of the MDFF Plugin within VMD.^40^ Five maps in total were generated for each state at varying resolutions from 1 Å to 9 Å increasing by 2 Å. These density-maps can be used directly without inversion to a grid potential; unlike MDFF, where density-maps need to be converted to grid-potentials to be incorporated as an energy term.^48^ The atoms selected to be coupled to the cryo-EM map depends on the map resolution. As a rule of thumb, based on *ab initio* electron density map refinements,^60,61^ data with resolutions between 4-8 Å are fitted to backbone atoms while resolutions higher than 4 Å are fitted to all protein atoms except hydrogen.

**Figure 2.**
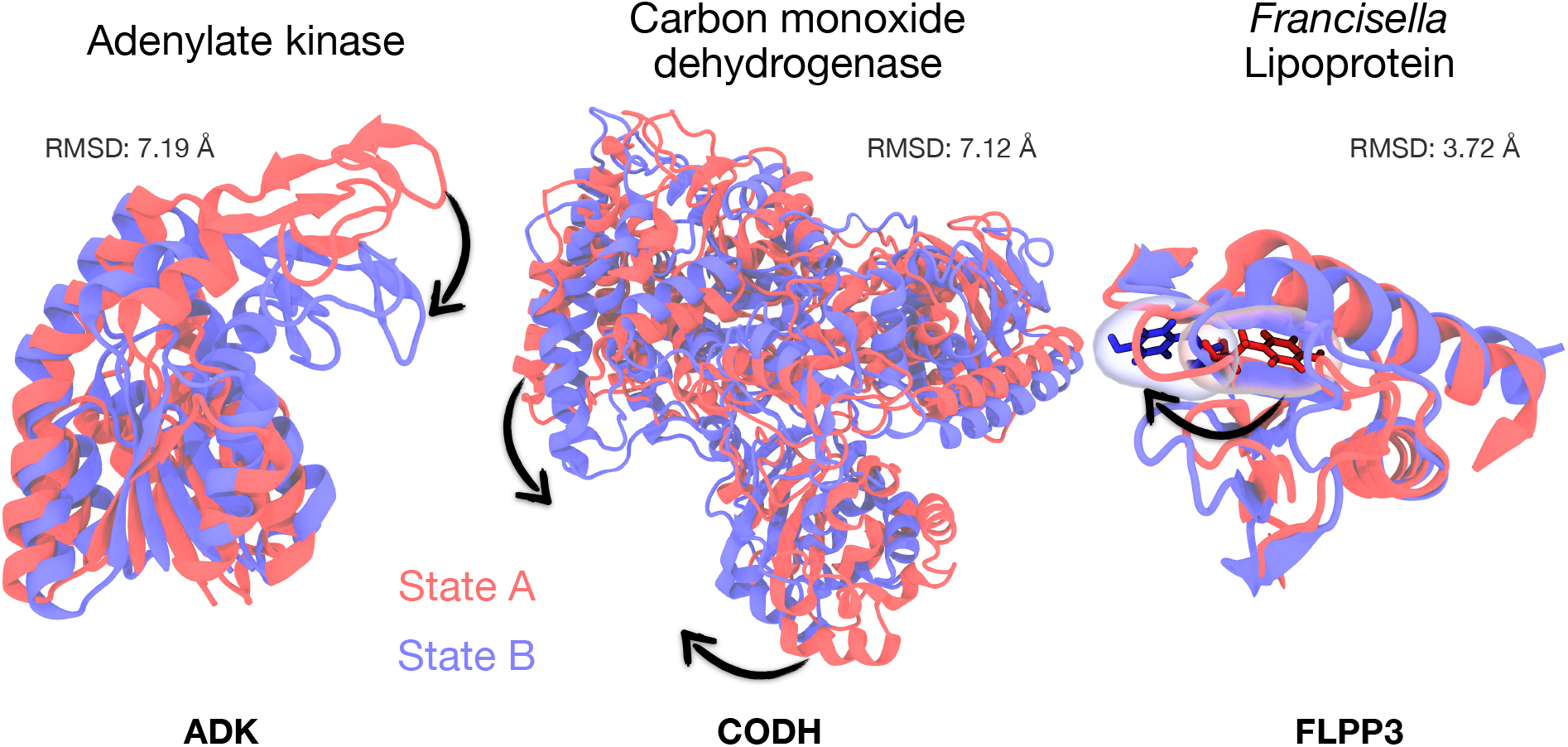
The graphical overlay of states A and B for three two-state protein systems used to study the Multi-Map collective variable. Adenylate kinase (ADK) states are defined by the open (A), PDBID 4AK^E53^ and closed (B), PDBID 1AKE^52^ states. The carbon monoxide dehydrogenase (CODH) A and B states are taken from chains D and C of the PDBID 1OAO^62^ structure. The FLPP3 A and B states are drawn from the crystal structure PDBID 6PNY^63^ and the NMR structure PDBID 2MU4,^64^ respectively. The flipped tyrosine residue, Tyr83, is highlighted within the FLPP3 structure.

Analogous to MDFF, biasing with the Multi-Map colvars requires secondary structure constraints utilizing the extraBonds feature in NAMD.^47^ These constraints, which retain secondary structure folds, prevent overfitting of the models to the maps. Additionally, positional and orientational constraints are used to confine the sampling in regions between adjacent cryo-EM maps, as discussed in the next subsection.

### D. Steered Molecular Dynamics (SMD)

The formulation above lays the groundwork for using the Multi-Map colvar to define a structure’s similarity to states defined by cryo-EM maps. Besides monitoring the configurational state of a protein, the Multi-Map colvar can be used to steer a protein configuration to a target map, *ϕ*. By employing a moving harmonic restraint, we can derive an initial pathway between states using atomic forces derived from the following equation.

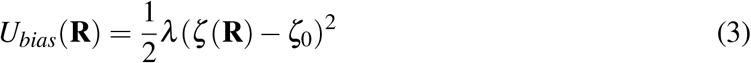

Here *λ* is the force constant, and *ζ*_0_ is the Multi-Map colvar target value, which changes uniformly over the course of the simulation. The force constant is chosen according to the range of colvar values required for describing a transition (Table S2). Expanding the harmonic and calculating the atomistic forces derived from *U_bias_*, one sees the force coming from the first term is proportional to *dϕ*/*dr_i_*. Similar to MDFF,^48^ this term localizes atoms onto the density surface. The second term, proportional to *d*(*ϕ*(**R**)*ϕ*_0_(**R**_0_))/*dr_i_*, is akin to taking the derivative of a correlation and acts to drive **R** → **R**_0_, effectively steering the atomic structures towards the target density *ϕ*_0_.

The quantity and quality of cryo-EM data are typically insufficient to refine atomic models with a high degree of accuracy using *ζ_AB_*-steered molecular dynamics. Therefore, we supplement the energy function *U_bias_* with additional constraints, notably *U_ff_*, the CHARMM all-atom additive potential as well as additional constraining terms to confine sampling. The following equation thus governs the simulation dynamics

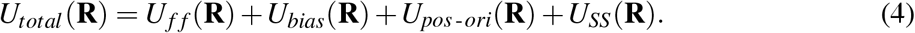

Here *U_SS_* is secondary structure restraints defined using the extraBonds feature of NAMD,^47^ and *U_pos-ori_* include center-of-mass and orientational restraints provided by the Colvars module.^37^ *U_SS_* maintains the set of secondary structure folds the systems starts with, and are typical for MDFF structure refinement.^49^ The *U_SS_* constraint could be omitted when folding from a random coil based on cryo-EM map data;^65^ however, the protein folding problem remains intractable in MD. Thus we start with a model that has commensurate folds to the target states. The term *U_pos-ori_* ensures that the system does not translate or rotate relative to the cryo-EM maps when biasing *ζ_AB_* near 0. Such treatment reduces the orthogonal degrees of freedom that do not contribute meaningfully to the transition pathway and is commonly seen in free energy perturbation simulations.^66^ The positional and orientational constraints ensure that structures derived along the pathway are relevant to the states defined by the cryo-EM maps.

### E. Bias Exchange Umbrella Sampling

Enhanced sampling methods are used to calculate the free energy change between two states in molecular simulations. Some of the well-established methods are umbrella sampling (US),^67^ adaptive biasing force method (ABF),^68^ metadynamics.^69^ We use an exchanging US algorithm to reconstruct the PMF along the reaction coordinate path sampled using SMD. Here, we briefly describe the method and application to the recently introduced system-specific reaction coordinate, cryo-EM map density collective variable, *ζ*(**R**). For biomolecular systems with large degrees of freedom, the sampling efficiency of US is significantly improved when combined with a replica-exchange scheme, hence the term bias-exchange (or replica-exchange) umbrella sampling.^70,71^ In replica-exchange MD or bias exchange umbrella sampling (BEUS), each replica (or window) is assigned a different value of a given property for the system. Periodic attempts are made to exchange between replicas using a rule defined by the Metropolis criteria. The exchange rule is set based on biasing potentials, attempting a swap every 500 steps (or 0.5 ps) over a range of 126 windows. The mixing of replicas in BEUS, ensures continuous sampling for protein conformations between each replica, generating a more reliable free energy profile for the process. To remove any unphysical bias towards a particular state, 50% of the windows i.e., 63 of them were initialized with models picked from the A→B SMD, evenly interspersed with initial models for the other 63 windows chosen from steering along the B→A direction. Further details on BEUS and applications to different biomolecular systems are discussed elsewhere.^58,70,72^

### F. Calculating Potentials of Mean Force

There are various methods of assessing the potential of mean force. For the steered molecular dynamics trajectories, the non-equilibrium work is computed internally by the Colvars module.^73^ The non-equilibrium work permits an initial free energy estimate based on the second law of thermodynamics, which has as a consequence that the work for a non-equilibrium process W is bounded from below by the overall free energy difference, *W* ≥ Δ*F*,^74^ although the short simulations typically substantially overestimate the difference. From the bias exchange umbrella sampling simulations, we estimate the free energy profiles and their uncertainties along the defined reaction coordinate using a modified version of BayesWHAM.^75^ The implementation has been accelerated by using Habeck’s Gibbs sampling method^76^ rather than Metropolis-Hastings sampling as originally implemented.^75^ As an additional check, multistate Bennett’s acceptance ratio calculations,^77^ as implemented in pyMBAR, are used to verify our methodology. Uncertainty estimates are obtained by trajectory subsampling to compute the variation in the computed free energies, which is used to assess convergence.

## III. RESULTS AND DISCUSSION

The changes in the Multi-Map colvar were monitored and assessed during equilibrium, nonequilibrium, and free energy simulations. We focus on how these colvars track global and local conformational rearrangements. The results bring to light the pros and cons of applying this reduced representation over traditional geometric collective variables emerging from a linear combination of atomic coordinates.

### A. Multi-Map colvars describe large scale structural transitions at > 5 Å density resolution

First, we seek to determine whether the colvar can distinguish between states A and B based exclusively on the *ζ_AB_* value computed from Eq. 2. Fig. 3A shows histograms of the *ζ_AB_* colvar distribution during an equilibrium simulation. At resolutions of 3-9 Å, we find that the conformations initiated in state A retain the negative values they start with, and similarly, configurations that start in state B sample around the positive values for the *ζ_AB_* coordinate. The initial values of the *ζ_AB_* colvar for simulations starting in either state A or B reflect a perfect fit to the data by construction and are near unity in the scaled *ζ_AB_* space. These are the minimum and maximum values the *ζ_AB_* colvar can assume. For the colvars derived from high-resolution density maps, the equilibrium conformations rapidly drift away from these limiting *ζ_AB_* values (Fig. S1). The *ζ_AB_* relaxes to distributions with a near-zero mean, implying minimal separation between the states, and is shown clearly through map correlation coefficient becoming equivalent (Figs. S4–S6). This trend of *ζ_AB_* approaching 0 observed with all high-resolution maps also suggests the need for sampling an intractably high number of conformations for estimating averages, irrespective of the system-size. In contrast, the degeneracy in conformations underlying a chosen *ζ_AB_* value is higher for the colvars derived at a lower resolution. This is reflected in the broader distribution of low-resolution *ζ_AB_* values, peaked about non-zero means (Fig. 3). The low-resolution *ζ_AB_* distributions clearly distinguish between states A and B, as indicated by the peak separation in Fig. 3B; even in cases where RMSD comparisons cannot distinguish between states because they are equally far from either starting configuration (Fig. S2).

**Figure 3.**
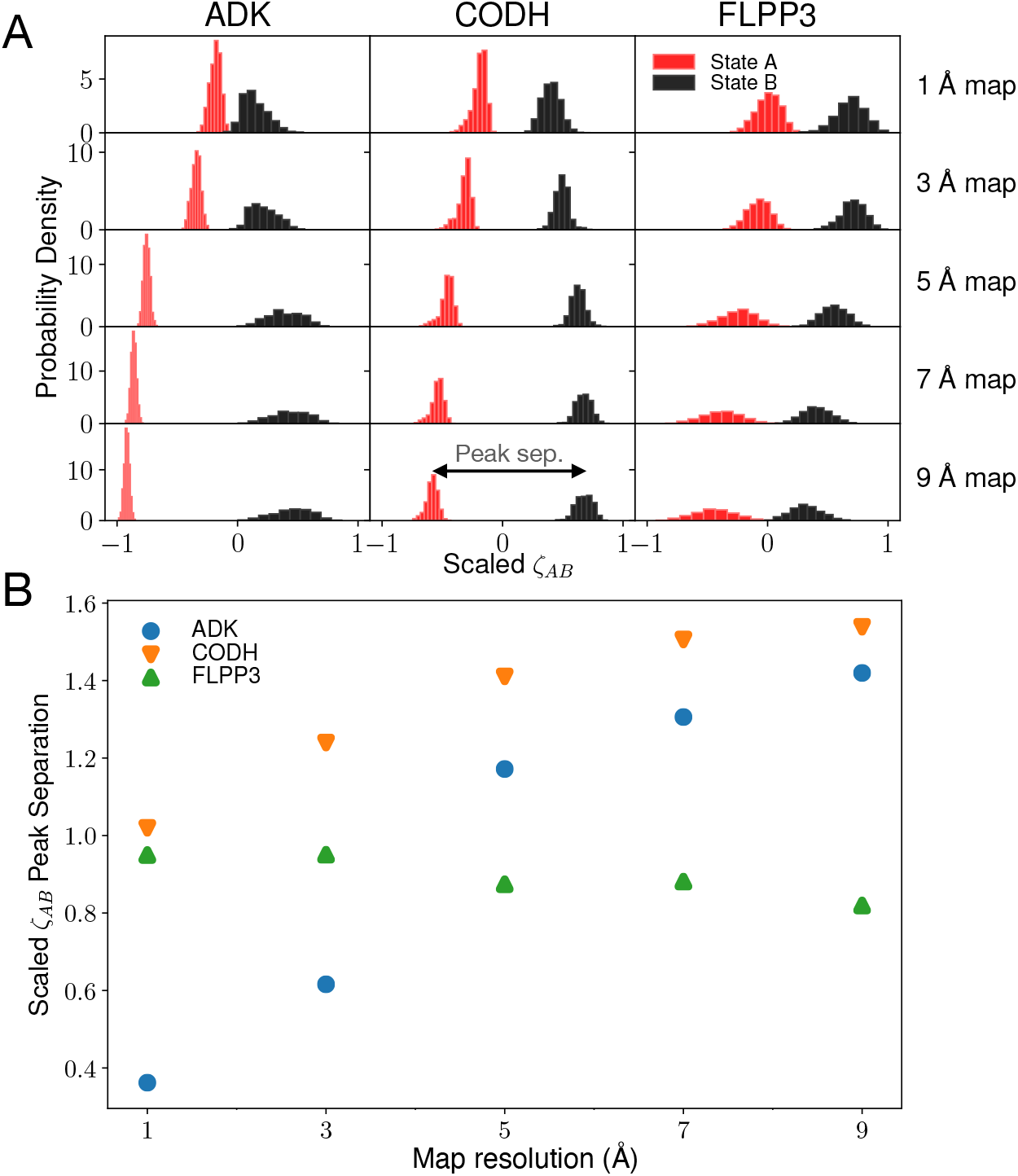
(**A**) Histograms of scaled *ζ_AB_* values from equilibrium simulations of ADK, CODH, and FLPP3 starting in either state A or B (red and black, respectively). The 10 ns following a 10 ns initialization period was used to construct the histograms (see Fig. S1 for the full equilibrium trajectories). The *ζ_AB_* values were normalized between −1 and 1 by dividing each value by half of the range seen during the equilibrium simulation. The double-sided arrow seen in the CODH 9 Å map plot shows a visual interpretation of peak separation. (**B**) Histogram maximum probability peak separation as a function of the map resolution used to derive the *ζ_AB_* colvar. The peak separation values are based on the scaled *ζ_AB_* profile, where the maximum separation that can be obtained is 2.

At any given resolution, the range of *ζ_AB_* values visited is the highest for CODH, followed by ADK and then FLPP3. This follows a trend guided by the number of atoms in these systems, where the range increases with increasing system size (Table S2). The separation between *ζ_AB_*(**R**_*A*_) and *ζ_AB_*(**R**_*B*_) is, therefore, most prominent in CODH and least in FLPP3. Conversely, for any system size, the range of *ζ_AB_* values decreases and finally plateaus with decreasing map resolution (Fig. 3B). Fuzzier density features for the maps of lower resolution have reduced values of *ϕ* for any *r_i_*, resulting in lower values of the *ζ_AB_* summation in Eq. 2. Despite this lower range of *ζ_AB_*, the separation of states improves dramatically at lower resolutions. For the 1 and 3 Å maps, *ζ_AB_*(**R**_*A*_) ~ *ζ_AB_*(**R**_*B*_) ~ 0. This effect is seen in Fig. S1, where the initial *ζ_AB_* colvar value moves quickly towards zero within the first 10 ns, and map correlations decay the fastest (Figs. S4–S6). For resolutions 5 Å or lower, the colvar tracks distinct large scale conformational changes, clearly representing the open and close states in ADK and CODH (Fig. 3A). At these resolutions, the separation between states on the *ζ_AB_* profile is roughly equivalent or higher to the corresponding RMSDAB scaled peak separation (Figs. 3B and S3 and Sect. S1) This finding suggests that the path length in *ζ* space is longer than or equivalent to the path created in the space of geometric collective variables composed of a linear combination of atom coordinates. The path through the extended space accessible to *ζ* promises exhaustive conformational sampling and the discovery of conformations hidden to the geometric collective variables. A more important benefit of *ζ* over the traditional geometric collective variables is that the knowledge of the endpoint structures is not required. Unlike RMSD-like variables where atomic models or structures need to be fit to each of the *K* maps to define a pathway, following Appendix A, *ζ* requires only the knowledge of one endpoint; obviating the need for *a priori* real space refinements, although the resolution needs to be accounted for (Sect. S2).

Molecular rearrangements for both ADK and CODH require large domain movements and have significantly larger RMSDs when compared to FLPP3 (Fig. 2). The FLPP3 conformational transition involves breaking an interior hydrogen bond made by Tyr83 and the tyrosine residue’s movement to an outward-facing conformation. The subtle rearrangements involved in the Tyr83 flip are only distinguishable with high-resolution maps (i.e., 1 and 3 Å). The lower resolution maps, and therefore the *ζ_AB_* colvar, cannot distinguish between FLPP3 configurations in state A or B. At low-resolution the cryo-EM maps for FLPP3 states A and B are highly similar with a correlation coefficient of 0.85, 0.92 and 0.95 for map resolutions 5, 7 and 9Å, respectively (Figs. S1 S6).

The *ζ_AB_* colvar has two requirements to be able to discern protein configurations into individual states. First, the transition between states needs to be large enough to distinguish between cryo-EM maps at nominal resolutions. Second, the cryo-EM map needs to be at a low enough resolution to incorporate an ensemble of structures undergoing thermal motion into a single state defined by the map. This first requirement is system dependent. The second requirement can be met by low-pass filtering of high-resolution cryo-EM maps using VMD’s voltools plugin if required.^31^ The blurring adds Gaussian halfwidths *σ* to the maps and enables the maps to incorporate more structures in their state definitions and thus a colvar which is better able to distinguish between the structural ensembles from states A and B.

### B. Steering MD along low-resolution Multi-Map variables produces complete transitions

Trial pathways for probing large scale conformational transitions are often generated using external forces via steered MD (SMD) simulations.^78,79^ Starting from states A or B for each of the three proteins, SMD was used to drive the transition to the other endpoint using the Multi-Map colvar defined at density map resolutions between 1 to 9 Å. The transitions were monitored by RMSD to both initial and final states (Figs. S8 and S9), the *ζ_AB_* components themselves (Fig. S10), and the density map correlations coefficients to both the initial and final states (Figs. 4 and S11). Regardless of the preferred metric, the steered trajectories consistently demonstrate that our enforced biases allow the initial state to approach the final state. At resolutions 5 Å or lower, the colvar traces for the forward and reverse paths derived from two independent simulations, strongly overlap. This result offers preliminary evidence of minimal hysteresis in the SMD protocol we used with *ζ_AB_* to induce A→B and B→A transitions.

**Figure 4.**
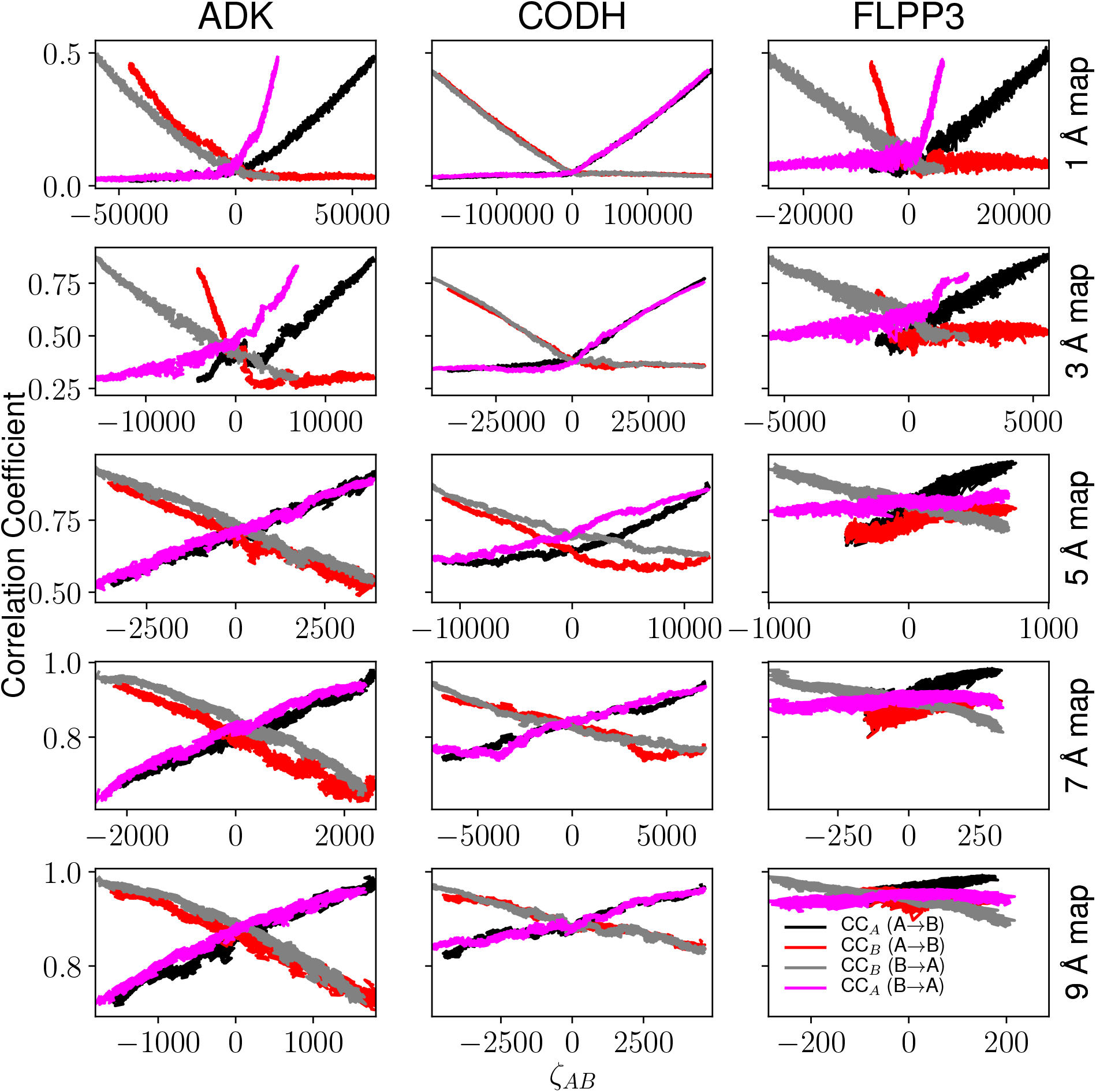
Traces of steered molecular dynamics trajectories highlight the relationship between *ζ_AB_* as defined in Eq. 2, and the correlation coefficient to the respective maps. Each subplot simultaneously shows results for the A→B (black and red lines) and B→A (gray and fuchsia lines) transition. Colored lines highlight motion towards the target, as measured through the cross-correlation to the target density, while the black and gray lines represent the falling out of the density.

At high-resolution (1 & 3 Å), ADK and FLPP3 are driven through transitions that appear to be complete along the correlation coefficient dimension but struggle to reach the extreme *ζ_AB_* values that are expected for states A or B. Incomplete transitions are due in large part to the well-defined local density features. At high-resolution, 1 & 3 Å, the density features are quite narrow and include sidechain conformations. Thermal motion prevents simulated systems from perfectly fitting to the maps, similar to the fast correlation decay observed in equilibrium simulation (Figs. S4–S5). Thus, for the high-resolution maps, the collective variable cannot find configurations that perfectly fit the maps and complete the transition in *ζ_AB_* space, even though more coarse-grained geometric collective variable metrics such as RMSD (Fig. S8) or density correlation coefficient (Fig. S11) indicate the transition has completed.

Low-resolution maps are also not without their issues within a steered simulation context. As visualized in Fig. 2 and noted in discussions of prior simulations,^63^ the transition in FLPP3 depends on the rotation of Tyr83 from packing in the interior to becoming solvent-exposed. This relatively subtle shift is difficult to capture in the context of low-resolution electron densities, unlike the much larger conformational changes for ADK and CODH (Fig. 2). For low-resolution maps, similar states have largely overlapping electron densities, resulting in comparatively few density differences the Multi-Map collective variable can exploit to drive conformational change. This is particularly clear in the variation of x-axis ranges in Fig. 4, where *ζ_AB_* varies less for FLPP3 along the transition than it does for the other systems tested. In summary, the small structural change for FLPP3 complicates transitions driven by low-resolution structural data, while the narrow densities for high-resolution data complicate the search process for poses that do not already fit the imposed density well. Nonetheless, with density data at ≈ 5 Å or lower resolution, large scale conformational transitions on the order of 7 Å change in RMSD are captured with minimal hysteresis between the forward and reverse pathways.

### C. Non-equilibrium work analysis reveals resolvability by flexible fitting

Driven transitions at the molecular level are typically non-equilibrium processes, accelerating slow transitions on the millisecond or longer timescale down to nanosecond timescales. Through analyzing non-equilibrium work, which tracks the accumulated forces and displacements along the SMD trajectory, it is possible to (i) estimate how resolvable a map is or “resolvability” of a map by flexible fitting,^39^ and (ii) glean the preferred sequence of events for a given transition.^80–82^ In Fig. 5, we analyze the non-equilibrium work from our own steered trajectories to evaluate the preferred directionality for these transitions.

**Figure 5.**
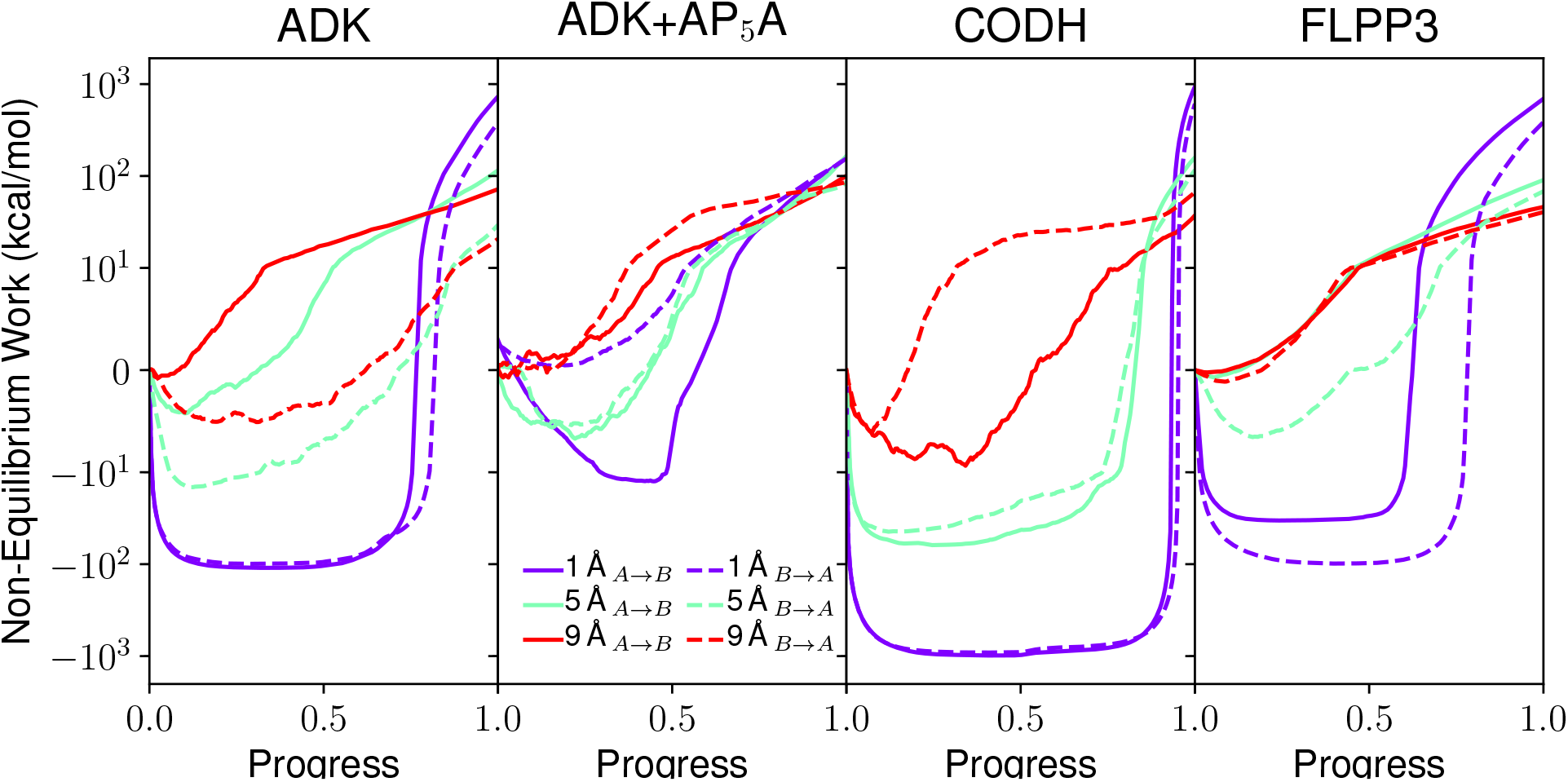
The non-equilibrium work for the SMD transitions in explicit solvent guided by either the 1, 5, or 9Å resolution maps. Solid lines indicate the A→B transitions, while dashed lines indicate B→A transitions. The map resolution is indicated by line color, with bluer colors indicating high-resolution maps, while redder colors indicate lower resolution maps. Note that due to the considerable variation in nonequilibrium work values, the plot is linear in the range (−10,10), and is plotted logarithmically outside of that range. Results for 3 and 7 Å resolutions are reported in Fig. S13.

SMD along the Multi-Map colvars indicates that there is an energy cost to flexible fitting. While the lower resolution maps require less work to fit a broad set of correct models into the map, higher resolution maps require more work to fit the correct model, as the number of such models non-linearly decreases with increase in resolution.^83^ The work needed for the fitting of higher resolution maps further increases with system-size, from FLPP3 to ADK to CODH. Establishing a common theoretical underpinning of why real-space refinement becomes more cumbersome for high-resolution maps,^84^ the non-equilibrium work profiles of the Multi-Map colvar explains several refinement challenges faced by MDFF, ROSETTA-EM or other density-guided MD protocols when the low-resolution EM refinement tools were originally re-purposed to resolve high-resolution density maps.^31,85^ A large amount of work is wasted to bring structures in the proximity of the density features when maps A and B are non-overlapping. This scenario is prominent with sub-3 Å density maps, showing accumulation of negative work, and implying that an automated MD refinement starting with arbitrary search models e.g., 3 Å RMSD away from the target model will waste 10 kcal mol^−1^ of energy before the flexible fitting to the density begins. Such physical limitations have proved detrimental in extending straight-forward MD refinements of maps between 3–5 Å resolution.^86^ At lower resolution, when the overlap between maps A and B improves, less work is wasted, and most of the MD is productive in fitting the model to the map. The refinement of structures is seen in their growth of correlation coefficients during the SMD (Figs. 4 and S12 and Table S2), and improved Molprobity^87^ scores for the associated structures (Table S3).

Leveraging the non-equilibrium work profile in Fig. 5, we can hypothesize the basic outlines of the free energy landscape for each system. For instance, for ADK without ligand, the B→A transition requires less non-equilibrium work than the A→B transition, suggesting that the *apo* state of ADK would prefer to be in state A (the open state) to accommodate a ligand. Conversely, when ADK is ligand-bound, the A→B transition requires marginally less work than the B→A transition, suggesting that the closed state B is slightly more probable than the open state A. The ligand-dependent energy surface is a well-known feature for ADK, and is qualitatively in agreement with previous findings.^88–90^ A similar analysis indicates that the A state would be favored in both CODH and FLPP3. For FLPP3 specifically, where the NMR-derived open state represented by state B is known to be more prevalent,^63^ this is further evidence that this reaction coordinate is not reliable in capturing relatively modest structural rearrangements.

### D. Free energy profiles with Multi-Map variables resolve large-scale conformational transitions

We test how a reaction coordinate defined by the difference between the two maps estimates the relative free energy differences between the two end states. To this end, we implement the BEUS protocols to derive free-energy differences along the *ζ_AB_* profile for the three examples. The windows are linearly distributed along *ζ_AB_* pathways of minimal hysteresis derived from Fig. 4.

#### 1. ADK open-to-close transition

To examine the relationship between free energy differences and map resolution, BEUS was run with *ζ_AB_* at five different map resolutions for ADK (see Fig. 6 left). At the highest resolutions between 1 to 3 Å the endpoints are thermodynamically inaccessible, consistent with the fact that only a handful of conformations can fit the distinct features of the high-resolution density maps. Thus, there is a frustration that entropically boosts free energy at extreme *ζ_AB_* values. These artificially sharp features on the energy landscape subside at the lower resolutions (≥5 Å, wherein the states close to the endpoints A and B (indicated by −1 and +1 values along the conformational coordinate) become more thermally accessible. There is disagreement in free energy differences between the high and low-resolution maps. While these differences are drastic for the 1 and 3 Å maps compared to the lower resolution maps, the 5, 7, and 9 Å maps are in agreement with each other in terms of both endpoint free energy values and their free energy differences between local minima.

**Figure 6.**
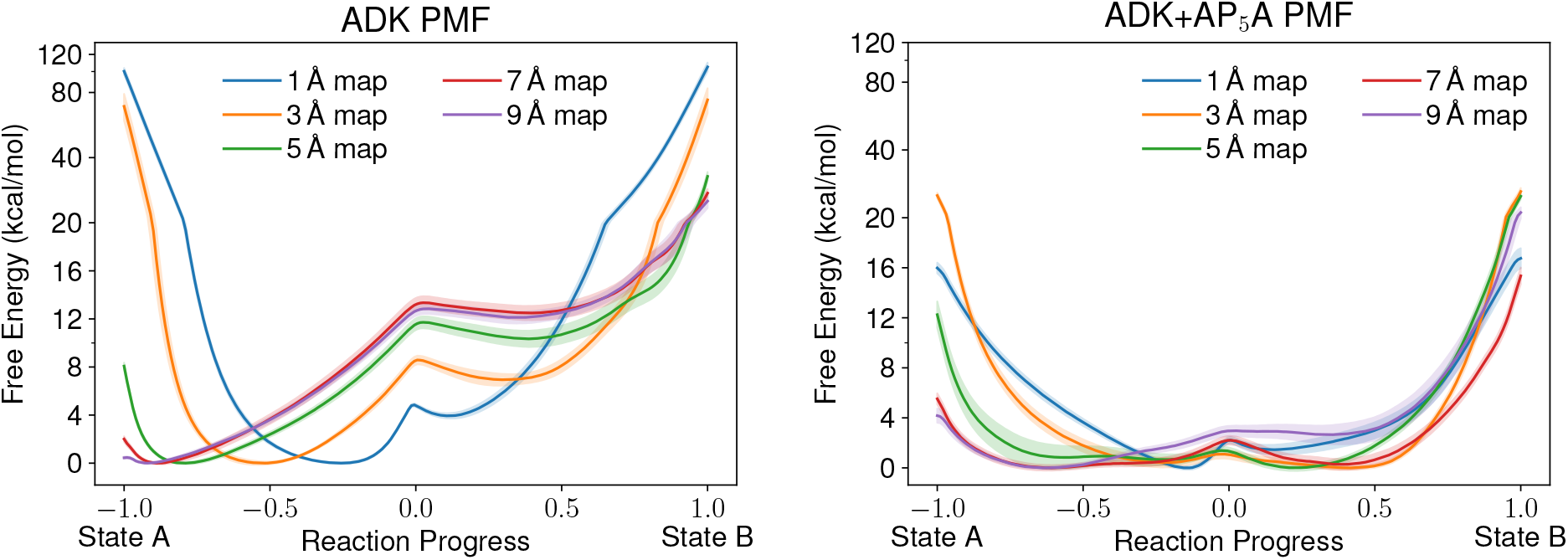
Free energy profiles for ADK without (left) and with (right) its AP_5_A ligand (right) using the Multi-Map reaction coordinate across tested resolutions. The x-axis coordinates have been scaled so that the reaction coordinate ranges are commensurate across different resolutions. Error estimates obtained by the spread of 5 ns trajectory subsamples are represented by the shaded regions around each free energy estimate. For convergence estimates based on subsamples, see Figs. S15–S24.

Fig. 6 shows free energy profiles for two ADK systems, apo and holo (AP_5_A inhibited). Free energy profiles of both ADK systems have been obtained in previous computational studies with high-resolution crystal structures.^88–90^ Notably, apo-ADK’s closed state (state B), is less favorable than apo-ADK’s open state (state A). Conversely, holo-ADK’s closed state is more favorable than the open state based on the literature.^88–90^ As shown in Fig. 6, the expected trends are retained, but now through the refinement of much lower resolution density maps rather than specific structures. Using the density maps of resolution 5Å or lower, apo-ADK’s free energy difference between states is approximately 10 kcal mol^−1^ which is comparable to the free energy differences found in previous studies.^88–90^ In contrast, the holo-ADK shifts the population towards the closed state B, with both the open and closed states having approximately equivalent free energy. Depending on the map resolution, AP_5_ A binding changes the well depths for both states to be within 1 kcal mol^−1^ of one another. The free energy trends also follow a reverse trend in vacuum, where the closed form is more stable than the open conformation by 16 kcal mol^−1^ (Fig. S14). Thus every time new interactions close the pocket in apo-ADK, be it through binding interactions with the AP_5_A ligand in the holo-state or via enhanced electrostatic interactions in vacuum, the closed conformation becomes more stable. Otherwise, apo-ADK is primarily open. This distribution of free energy implies that induced fit (and not conformational selection) is at work to enable AP_5_A binding to ADK, a result that matches kinetic assays and NMR results.^91,92^

#### 2. Sidechain flip in FLPP3

To elucidate the limitation of the Multi-Map colvar, we present the free energy profile for Tyr83 flip and the associated pocket opening in FLPP3 (Fig. 7). We have recently isolated the open, close and occluded conformations of FLPP3 using serial femtosecond crystallography (SFX) and NMR spectroscopy.^63,64^ Umbrella sampling simulations enabled the hierarchization of these conformations in terms of distinct Tyr83 orientations,^63^ whereby the open conformation with solvent-exposed Tyr83 was found to be significantly more stable. Already depicted in Fig. 4, the MultiMap colvar fails to capture the complete transition between the Tyr83-flipped end states of FLPP3. Unlike ADK, where the most separated end states were observed 5 Å or lower resolutions, the most resolved FLPP3 states are seen at 3.0 Å (Fig. 3). Under these conditions, the PMF reveals almost equally probable open and close states on either side of *ζ_AB_* equal to 0. At lower resolutions (≥5 Å), the converged statistics favor closed state over the open state. This result is in stark contrast to NMR that favors the open state to the close state by a population ratio of 2:1.^63^ The closed structure seen in the SFX data is over stabilized by lattice contacts, and is, therefore, even rarer. Thus, for this system, the BEUS of the Multi-Map colvar converged to unreliable results. This failure can be rationalized using Fig. 4. Unlike the ADK and CODH, the end states for FLPP3 are only accessible to the Multi-Map colvar at higher resolutions, *albeit* with hysteretic artifacts. Since the sidechain motions contribute minimally to the overall map transformation, both the solvent-exposed and buried Tyr83 were almost equally likely. Thus, the open and closed states of FLPP3 were found to be almost equi-energetic. At lower resolutions, the open →close transition appears more likely during SMD, with the reverse movement progressing only midway along the conformational coordinate. Occupancy of the buried Tyr83 within the lower resolution map is unphysically higher than that of the solvent-exposed orientation. Consequently, at resolutions of 5 Å and beyond, the open to close transition in FLPP3 is found to be exergonic. We expect a slower SMD to allow more chances for the Tyr83 to flip outward, and therefore address this error, possibly only at higher resolutions given the small-scale structural transition at hand.

**Figure 7.**
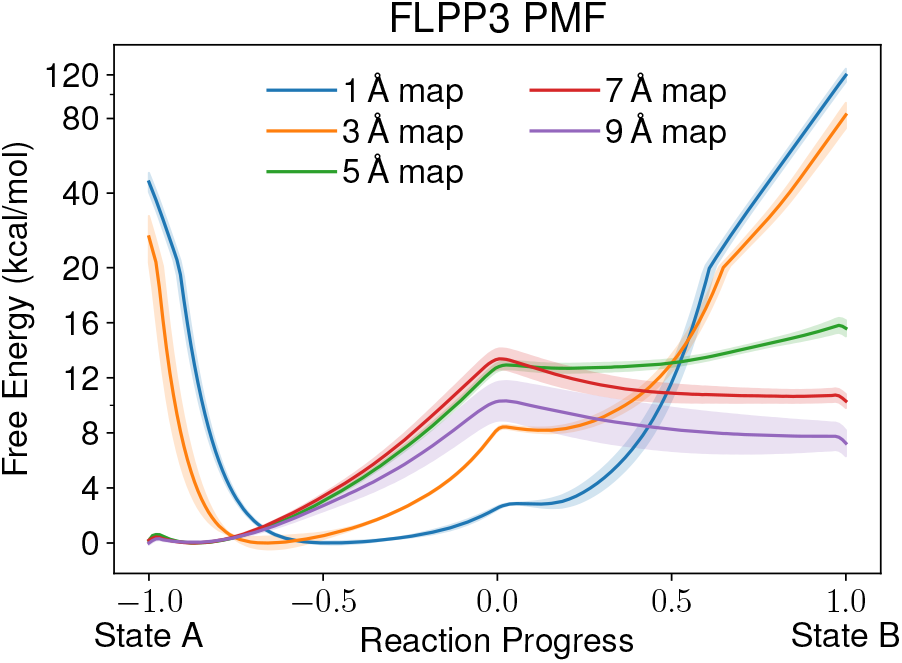
Free energy profiles for FLPP3 using the Multi-Map reaction coordinate across tested resolutions. The x-axis coordinates have been scaled so that the reaction coordinate ranges are commensurate across different resolutions. Error estimates obtained by the spread of 5 ns trajectory subsamples are represented by the shaded regions around each free energy estimate. For convergence estimates based on subsamples, see Figs. S30–S34.

#### 3. Hinge-bending in CODH

The largest system we applied the Multi-Map reaction coordinate to was CODH. Unlike PMFs for ADK and FLPP3, which were relatively consistent in shape between different input map resolutions, CODH demonstrates significant changes in the shape of the free energy profile in a resolution-dependent manner (Fig. 8). For similar entropic considerations mentioned previously, the PMF minima are near *ζ_AB_* =0 for 1 and 3 Å maps. The profiles for these high-resolution maps have sharp edges due to the difficulty of fitting thermalized states into a high-resolution map. The valley broadens when 5 Å resolution maps are applied to the Multi-Map colvar, offering comparable probability to the open and closed CODH structures, marginally skewed towards state A, while 7 and 9 Å maps are skewed towards state B.

**Figure 8.**
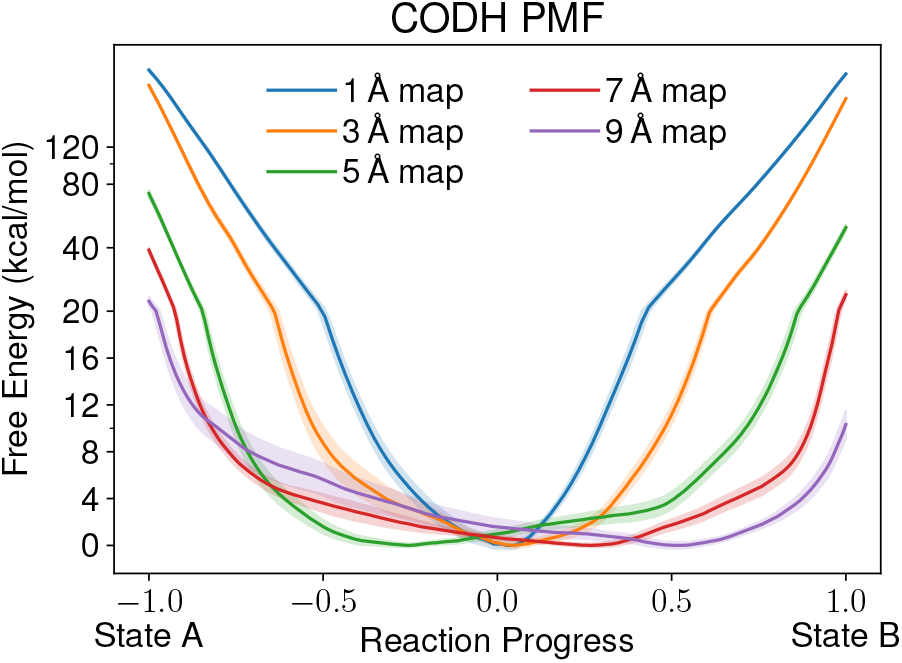
Free energy profiles for CODH using the Multi-Map reaction coordinate across tested resolutions. The x-axis coordinates have been scaled so that the reaction coordinate ranges are commensurate across different resolutions. Error estimates obtained by the spread of 5 ns trajectory subsamples are represented by the shaded regions around each free energy estimate. For convergence estimates based on subsamples, see Figs. S25–S29.

This energy landscape is consistent with the biochemical knowledge available for CODH, which implies that the protein structure fluctuates to accommodate substrate ingress.^93^ Indeed, the existence of both states within a single crystal structure implies that at least in crystallization conditions,^62^ the two states are equally probable and thus equal in free energy. This scenario is akin to classic conformational selection,^94^ highlighting that both conformations are thermally accessible in the absence of CODH substrates. Given the observation of substrate access tunnels in both states, while only state B is thought to be competent for chemistry,^62^ allosteric regulation for the CODH structure may be one mechanism for guiding metabolism through this multifunctional enzyme.^95^ Taken together, between the ADK and CODH examples the Multi-Map colvars capture two of the most universal mechanisms of allosteric interactions.

## IV. CONCLUSION

The use of volumetric maps as sources of external potentials in MD simulations has allowed the development of many enhanced sampling methods such as grid-steered MD,^51^ MDFF,^31,48^ and atom resolved Brownian dynamics.^96^ The most recent addition to these methods is the MultiMap variable,^37^ which is applied here to monitor transitions in protein structure based on electron density maps from cryo-EM or crystallography. Transformations between two protein states are successfully sampled for different protein sizes and types of conformational change, suggesting that the method is generalizable to a series of cryo-EM maps akin to the outcome of the manifold-based cryo-EM data analysis.^29,56^ In addition to the simultaneous utility of the Multi-Map colvar towards non-equilibrium work and free energy estimation, we here demonstrate its usefulness in the refinement of map-structure correlations coefficients and model-quality without requiring high-quality search structures. The Multi-Map formulation offers a statistical mechanical description for the resolvability of a map, addressing a point of concern in cryo-EM modeling.

Previous work demonstrated that the Multi-Map variable can characterize shape changes in supra-molecular aggregates such as biological membranes or confined-water pockets.^37,97^ The results shown here also open the door to its application together with a number of geometric or alchemical free energy methods focused on protein conformational cycles.^98^ Furthermore, in the specific application of structure refinement, data-driven approaches such as MELD^65^ or metainference^99^ can readily employ data simulated using the Multi-Map variable as a source of coarsegrained information for computing the Bayesian priors. From a biophysical standpoint, two of the most universal allosteric pathways, namely induced fit and conformational selection, were here successfully investigated. Though the free energy studies were found to be most efficient using low-resolution maps, the estimates are comparable to those determined from high-resolution structures.

## V. ACKNOWLEDGEMENTS

We acknowledge start-up funds from the SMS and CASD at Arizona State University. This research used resources and is authored in part by the Oak Ridge Leadership Computing Facility at the Oak Ridge National Laboratory, which is supported by the Office of Science of the U.S. Department of Energy under Contract No. DE-AC05-00OR22725. AS acknowledges CAREER award from NSF (MCB-1942763) and NIH/R01GM095583. JWV acknowledges the support by the National Science Foundation Graduate Research Fellowship under Grant No. 2020298734. The authors acknowledge Research Computing at Texas Advanced Computing Center (TACC) at The University of Texas at Austin and the Arizona State University for providing HPC, visualization, database, or grid resources that have contributed to the research results reported within this paper.

## VI. DATA AVAILABILITY

The data that supports the findings of this study are available within the article and its supplementary material.

## Appendix A: Estimating endpoints for the Multi-Map colvar for different resolutions

The Multi-Map collective variable used here is defined in compact form as *ζ_AB_* (Eq. 2) by using two 3-dimensional volumetric maps *ϕ_A_*(**r**) and *ϕ_B_*(**r**), which are synthetically generated but are treated otherwise as experimental data. Based only on the atomic density maps, it is thus possible to estimate what the colvar range should be. Given an atomic configuration **R**, a synthetic map *ϕ*_**R**_(**r**) may be generated and its cross-correlation with *ϕ_A_*(**r**) defined as:

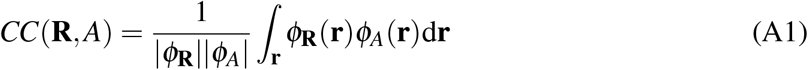

and similarly for *CC*(**R**, *B*). Inserting Eq. A1 into Eq 2, we arrive at

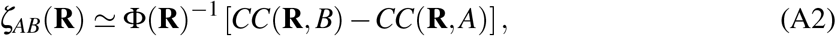

where the approximation lies in assuming that the density map *ϕ*_**R**_ is used in lieu of the precise atomic coordinates **R**. Also, Φ(**R**) = ∑*_i_w_i_ϕ*(**r**_*i*_) is the term of the Multi-Map variable evaluated at the coordinates **R** that best fit the set of local map *ϕ*. It is essential at this stage to recognize that correlation coefficient values fall in the range (0,1) due to non-negative densities. For instance, if a 0 Å resolution map were to exist, the densities would be delta functions with values equal to the atomic weight (*w_i_*) meaning that Φ(**R**)^−1^ = (∑*_i_w_i_*)^−1^. For other resolutions, Φ(**R**)^−1^ will similarly be a scaling factor related to the total atomic weight and will be invariant for well-fitted maps with equivalent resolution. Stated concretely for this special mass-conserving transition between states A to B,

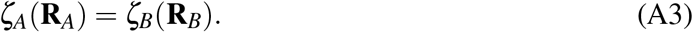

Assuming that a map is a Gaussian mixture model, the equality holds if both maps have homogenous and equivalent resolutions. If the resolutions are not equivalent then a scaling factor is needed to maintain the equality, as outlined below. However a more general boundary condition is

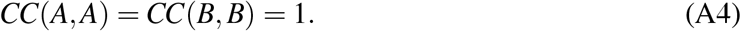

Combining Eqs. A2 and A3:

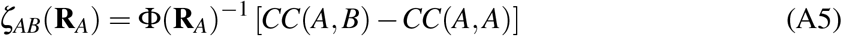

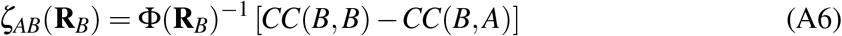

therefore:

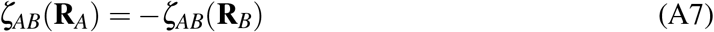

because *CC*(*A, B*) = *CC*(*B, A*). In practice Eq. A7 is limited by discretization errors, because both maps are interpolated onto a grid. However, by comparing the numerical and theoretical values 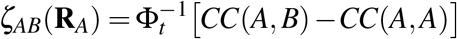 determined from Eqs. 2 and A2 in Table S2 within Sect. S2 for a range of resolutions, we find that a 1Å grid spacing has acceptable numerical error.

As a consequence of Eq. A7, the target value for *ζ_AB_* can be estimated, knowing only one endpoint from the transition for two maps with equal resolution. Importantly, all other atomic configurations other than **R**_*A*_ or **R**_*B*_ will generate *ζ_AB_* values whose magnitude is less than |*ζ_AB_*(**R**_*A*_)| = |*ζ_AB_*(**R**_*B*_)|, as the correlation coefficients are bounded and the density weight is conserved (Eq. A3).

If two maps have unequal resolutions with comparable local densities, the range for *ζ_AB_* will depend on the ratio between Φ_*A*_(**R**_*A*_) and Φ_*B*_(**R**_*B*_), like

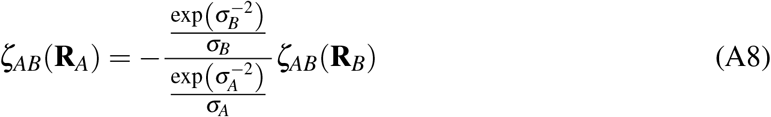

Where *σ_A_* and *σ_B_* are the resolutions of map A and B. The local density is calculated as the sum of atom weights divided by the map volume, which can be computed in VMD. Thus, even when the resolutions of the contributing maps are different, but their local densities are similar, we can still determine *ζ_AB_*(**R**_*B*_) with knowledge of only one high-quality structure across a series of maps. However, when the resolution of a map is nonuniform and the local densities do not match between the states A and B, the assumptions of Eq. A7 and A8 fail. In these cases, structural information on both **R**_*A*_ and **R**_*B*_ are needed *apriori* to determine the endpoint values of *ζ_AB_* for subsequent application in equilibrium and non-equilibrium MD or enhanced sampling simulations.

These considerations are non-trivially generalizable to capture conformational changes across the entire series of *K*-maps, which are considered a sum of 2-map transformations. Thus, by knowing the structure R at only one endpoint, the Multi-Map colvar allows in principle the construction of ensembles and simultaneous real-space refinement for each of the *K* maps contributing to the colvar. In practice, the variation of the local resolution within a cryo-EM map may prevent the application simple scaling rules to all the maps. Nonetheless, the nominal resolution of majority of the maps coming from conformational analysis with EM have highly comparable resolution as seen in the ribosome,^28^ RyR1 receptor,^29^ and recently in spike protein.^100^

## Appendix B: Implementation and Availability of the Multi-Map Collective Variable

The derivation and implementation of the Multi-Map variable are documented in ref.^37^. The implementation leverages recent improvements to the GridForces^51^ and Colvars^73^ modules, both of which are freely available in the most recent version of NAMD.^47^ Up-to-date documentation and input file fragments for several use cases are available at: https://colvars.github.io/colvars-refman-namd/colvars-refman-namd.html#sec:cvc_multimap

A NAMD configuration file fragment for invoking the two-state colvar (i.e. where the coefficients of the Multi-Map variable are −1 and +1, respectively) is provided below:

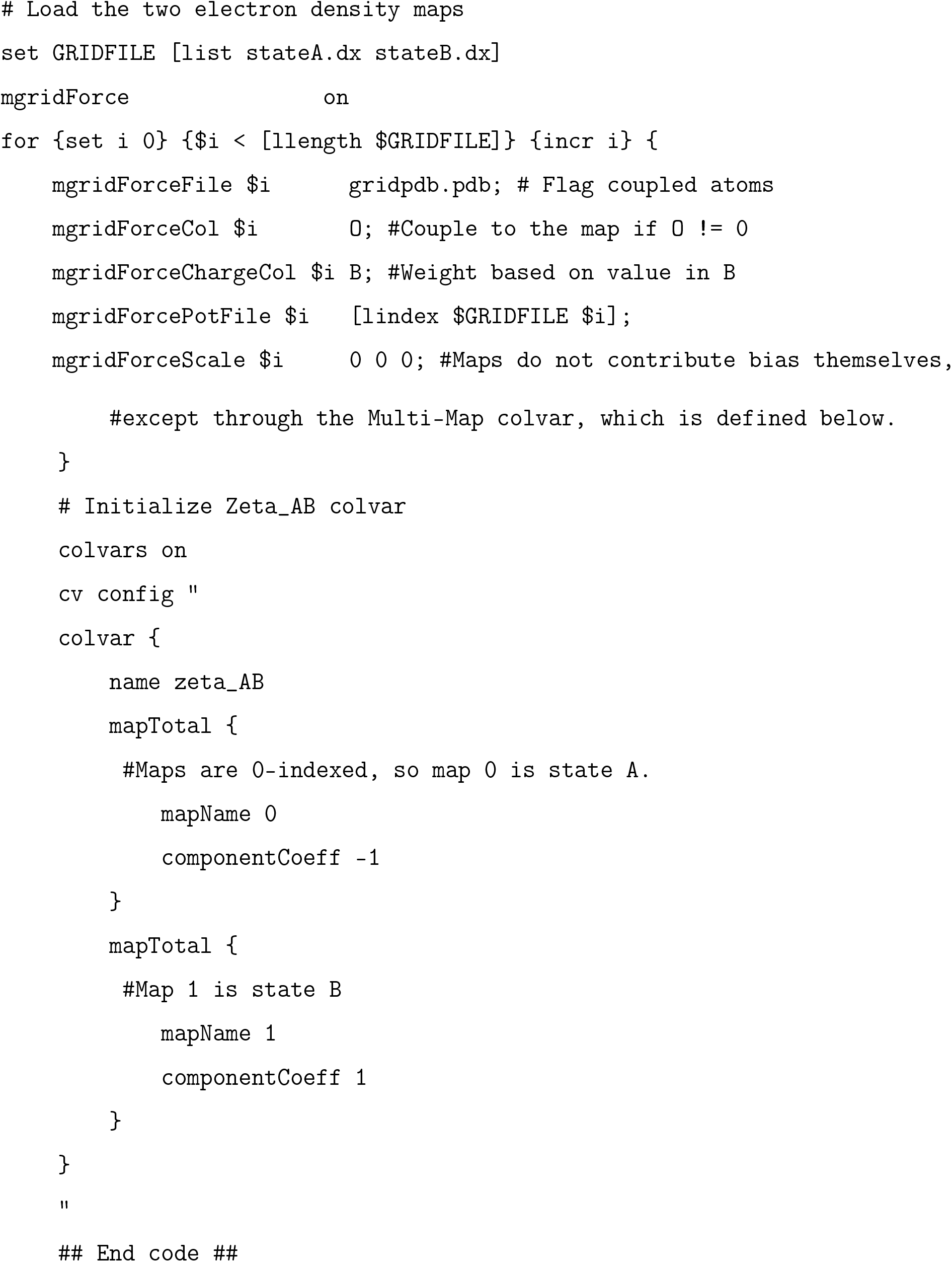

