## Supplementary material for "Data-guided Multi-Map variables for ensemble refinement of molecular movies": SI

(Dated: 13 October 2020)

### S1. CALCULATING RMSD$_{AB}$

The following variable was used to compare the separation of states achieved by the $\zeta_{AB}$ colvar on a scaled profile.

$$RMSD_{AB}(\mathbf{R}) = RMSD_{B}(\mathbf{R}) - RMSD_{A}(\mathbf{R}) \tag{S1}$$

Where RMSD$_A$ uses structure $\mathbf{R}_A$ as the reference. Likewise for RMSD$_B$. The value of $RMSD_{AB}(\mathbf{R})$ is thus positive for structure $(\mathbf{R}_A)$, and negative for $(\mathbf{R}_B)$.

Scaling was achieved by calculating a range based on the minimum and maximum values seen during equilibrium simulations starting in state A and B. All values were divided by the half of the range to yield profiles which lie between -1 and 1. Peak separation was calculated on the scaled profile for $\zeta_{AB}$ and RMSD$_{AB}$ by calculating the difference on the scaled profile between the most probable value for state A and B. Equilibrium values of $RMSD_A$ and $RMSD_B$ for both states and all systems are seen in Fig. S2.

### S2. PREDICTING MULTI-MAP COLVAR RANGE

Based on our numerical results, the range of $\zeta_{AB}$ depends heavily on system size, map resolution, and the CC between maps A and B. Incorporating these numerical results into a functional form dependent on the resolution $\sigma$, we observed data the following relationship between resolution and the maximum system-dependent $\zeta_{AB}$ value:

$$\zeta_{AB}^{max} = \left(\frac{m}{\sigma} + b\right)(1 - CC(A,B)) \sum_{i=1}^{N} w_i. \tag{S2}$$

Within Eq. S2, the variables $m$ and $b$ come from a linear fit across resolutions for a given system. For our systems, $m$ varies between 2.16 and 2.70, and $b$ ranges from 0.36 to 0.74. The corresponding predictions and their errors are presented in Table S2. Fig. S7 shows the linear fit to the data for each system. From the m and b values for each system, we see that the smaller systems, FLPP3 and ADK, scale similarly with resolution. While the larger system, CODH, shows a slightly different relationship with the resolution. This prescription is important for proteins, such as apoferritin, where multiple datasets are available in the EMDB at distinct resolutions. Fits such as Eq. (S2) can be made across a number of these resolutions to interpolate the limiting values of $\zeta_{AB}$ for all other resolutions, without the need to determine structures corresponding to every resolution.

TABLE S1. Molecular dynamics simulation parameters.[1–3]

| **Parameter** | **Vacuum** | **GBIS** | **Explicit** |
|---|---|---|---|
| Time Step (fs) | 2.0 | 2.0 | 2.0 |
| Cutoff (Å) | 12.0 | 16.0 | 12.0 |
| Switch (Å) | 10.0 | 15.0 | 10.0 |
| Temperature (K) | 300.0 | 300.0 | 300.0 |
| Pressure (bar) | NA | NA | 1.01325 |
| PME Grid Spacing (Å) | NA | NA | 1.0 |

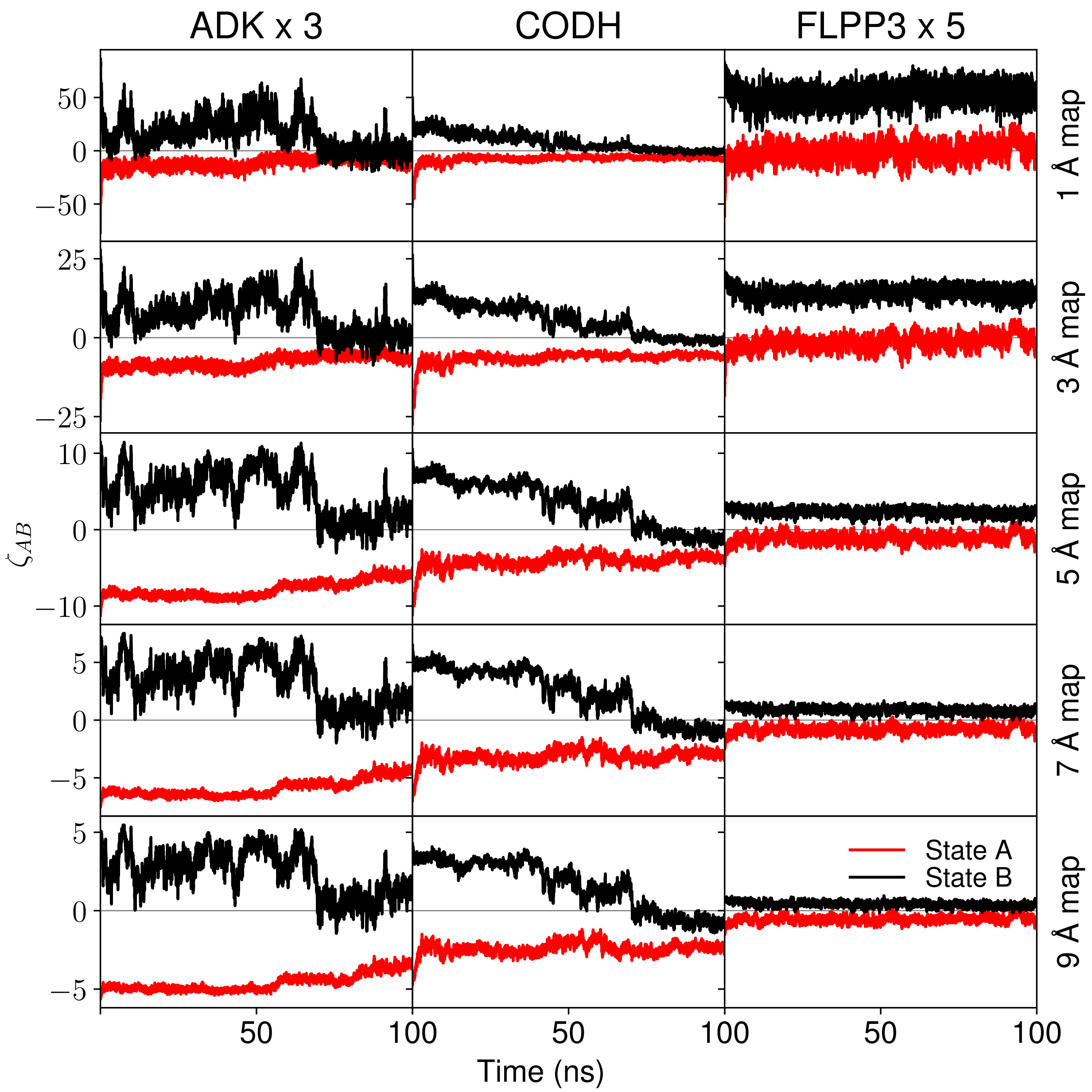

FIG. S1. The red and black lines show MultiMap colvar traces for the system starting in state A and B, respectively. Multi-map colvar values were monitored using maps with 5 different resolutions as indicated on the right. Due to the vastly different scales, the values for different systems are multiplied to maintain similar scales across rows for visualization.

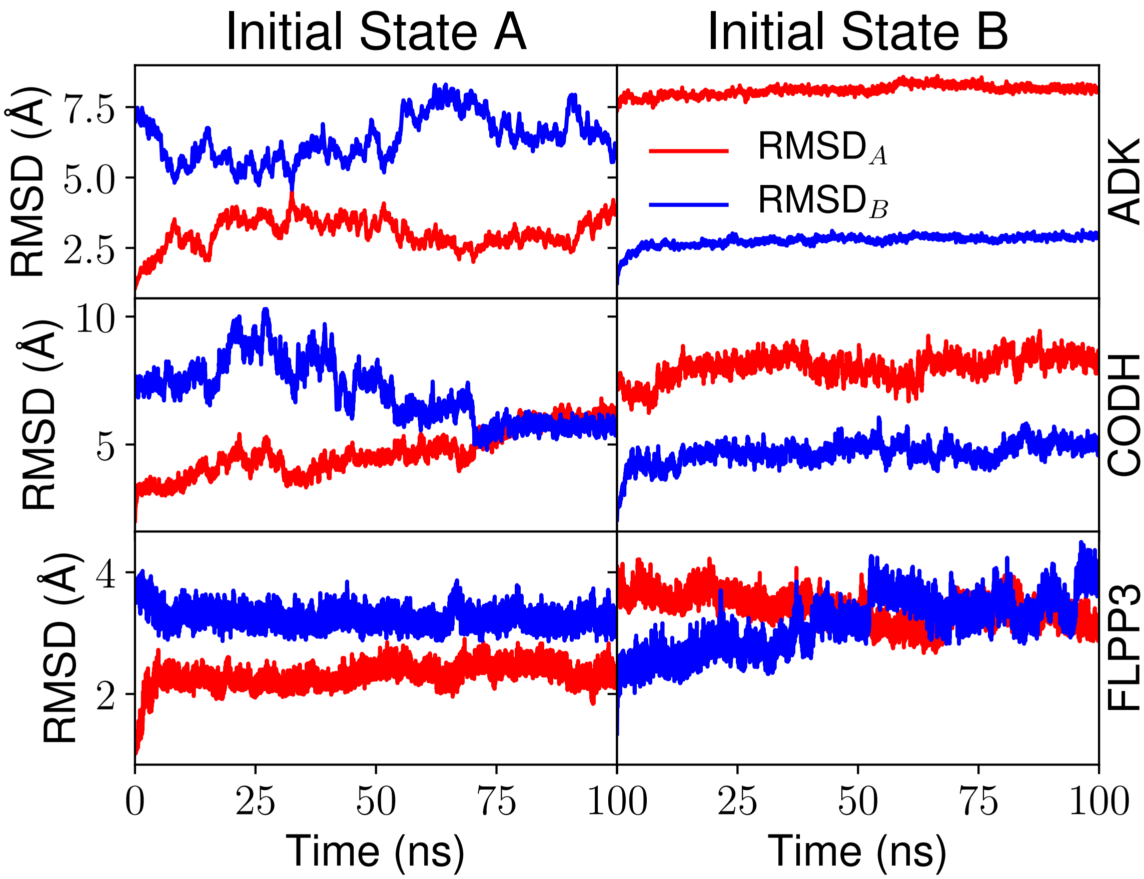

FIG. S2. RMSD traces for equilibrium trajectories initialized from state A (left) or B (right) for the three systems under study. The reference states for the traces are the initial structures for state A (red) or B (blue).

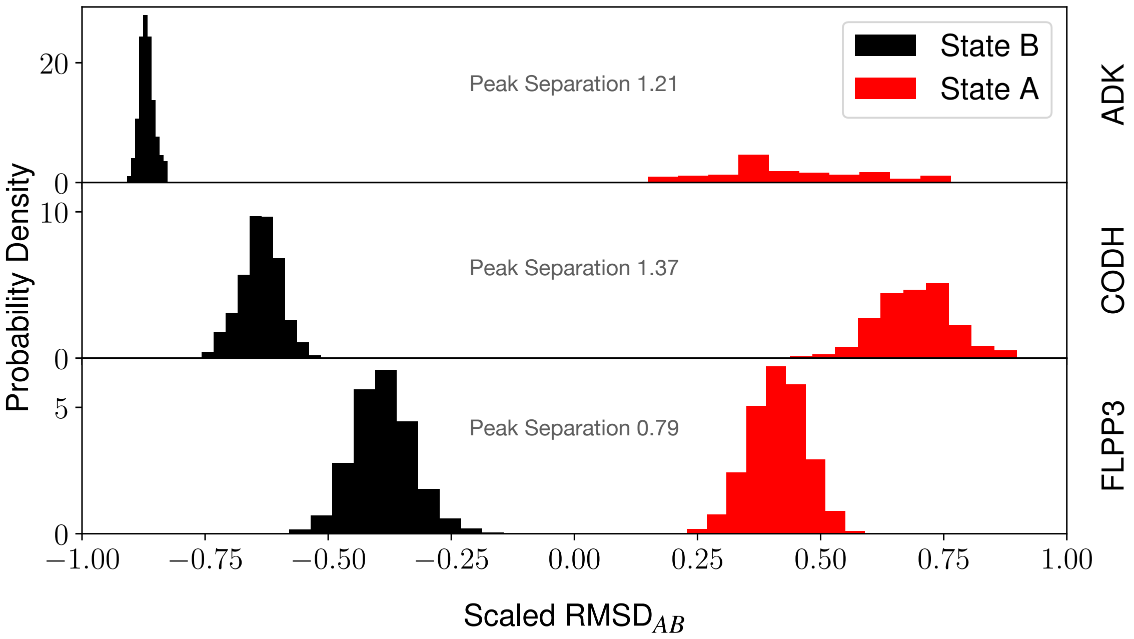

FIG. S3. RMSD histograms for equilibrium trajectories (10-20 ns) initialized from state A (red) or B (black) for the three systems under study. Scaling is acheived by dividing each value by half of the range seen durring the entire equilibrium trajectory. The reference states for the traces are the initial structures for state A or B. Peak separation is determined by taking the difference on the scaled RMSD$_{AB}$ axis between the most probable points.

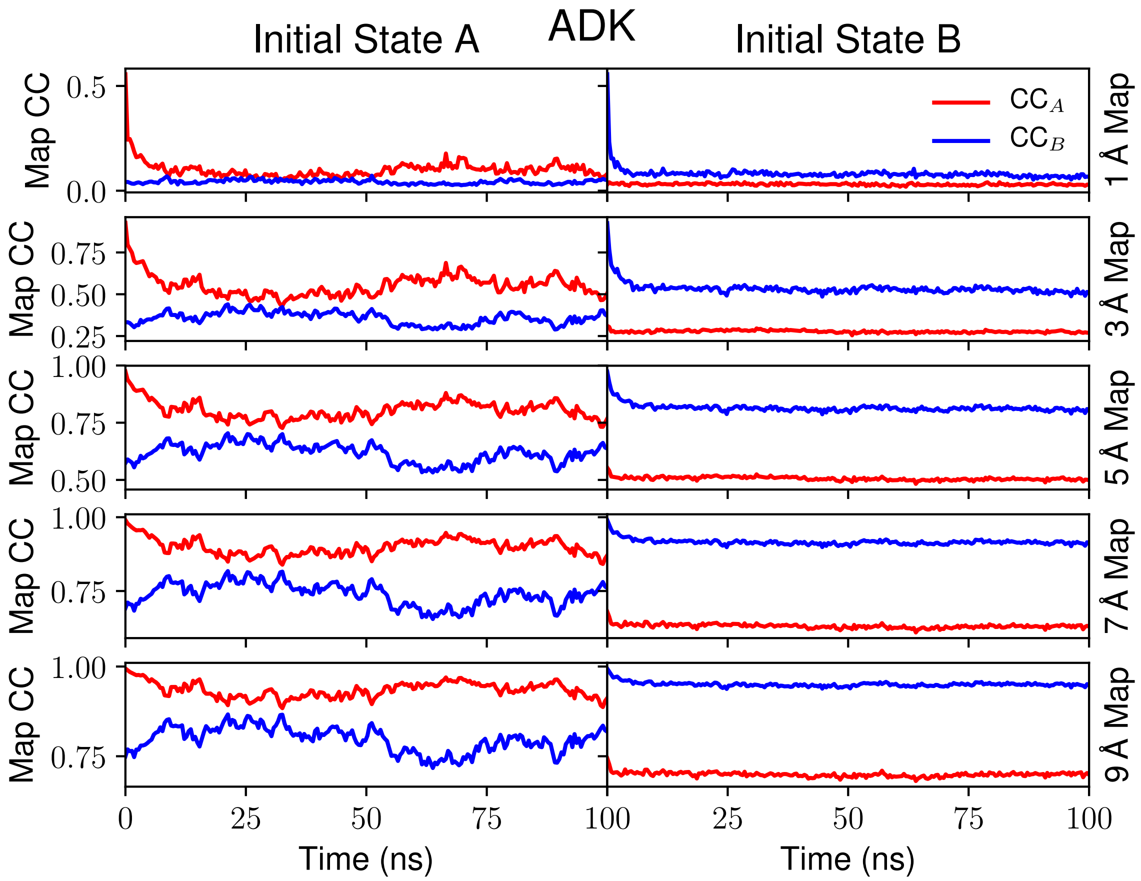

FIG. S4. Map cross correlation traces for equilibrium trajectories initialized from state A (left) or B (right) for ADK with different map resolutions. $CC_A$ is the cross correlation with respect to the state A map. Likewise, $CC_B$ is the cross correlation with respect to the state B map.

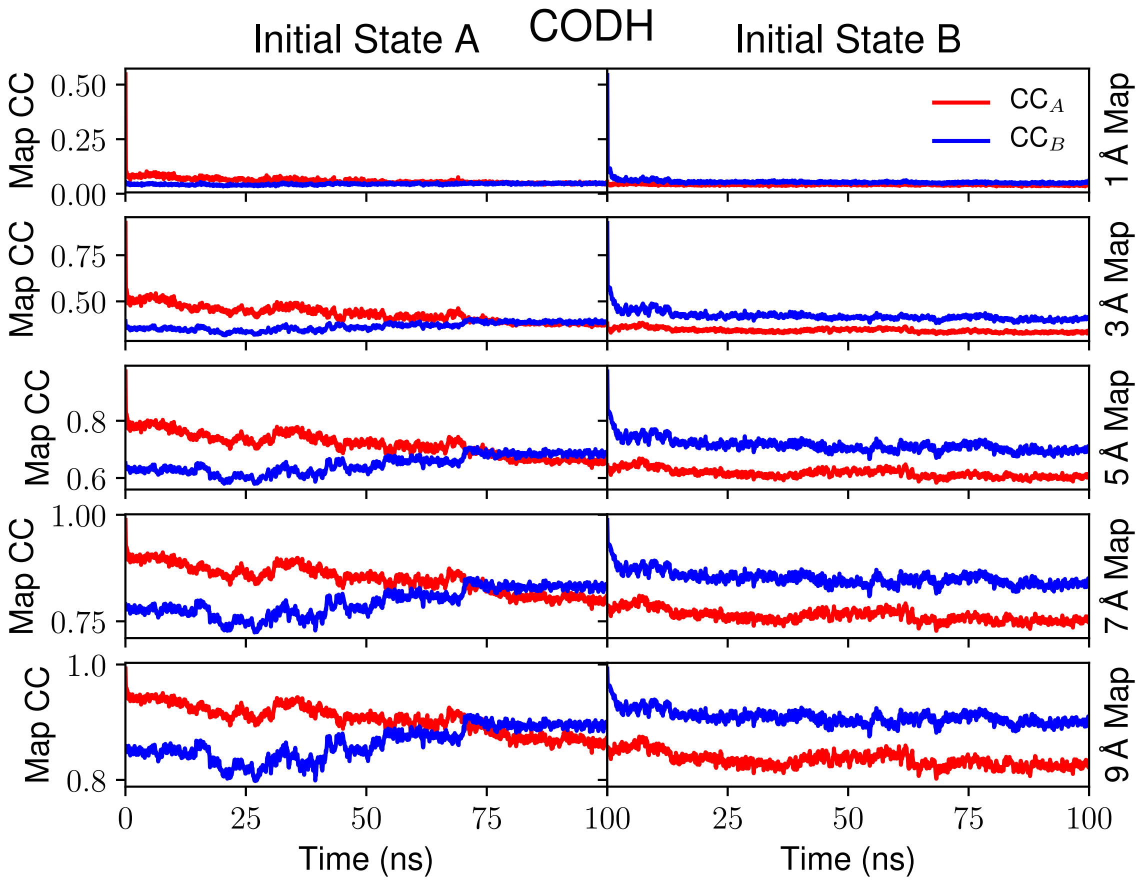

FIG. S5. Map cross correlation traces for equilibrium trajectories initialized from state A (left) or B (right) for CODH with different map resolutions. $CC_A$ is the cross correlation with respect to the state A map. Likewise, $CC_B$ is the cross correlation with respect to the state B map.

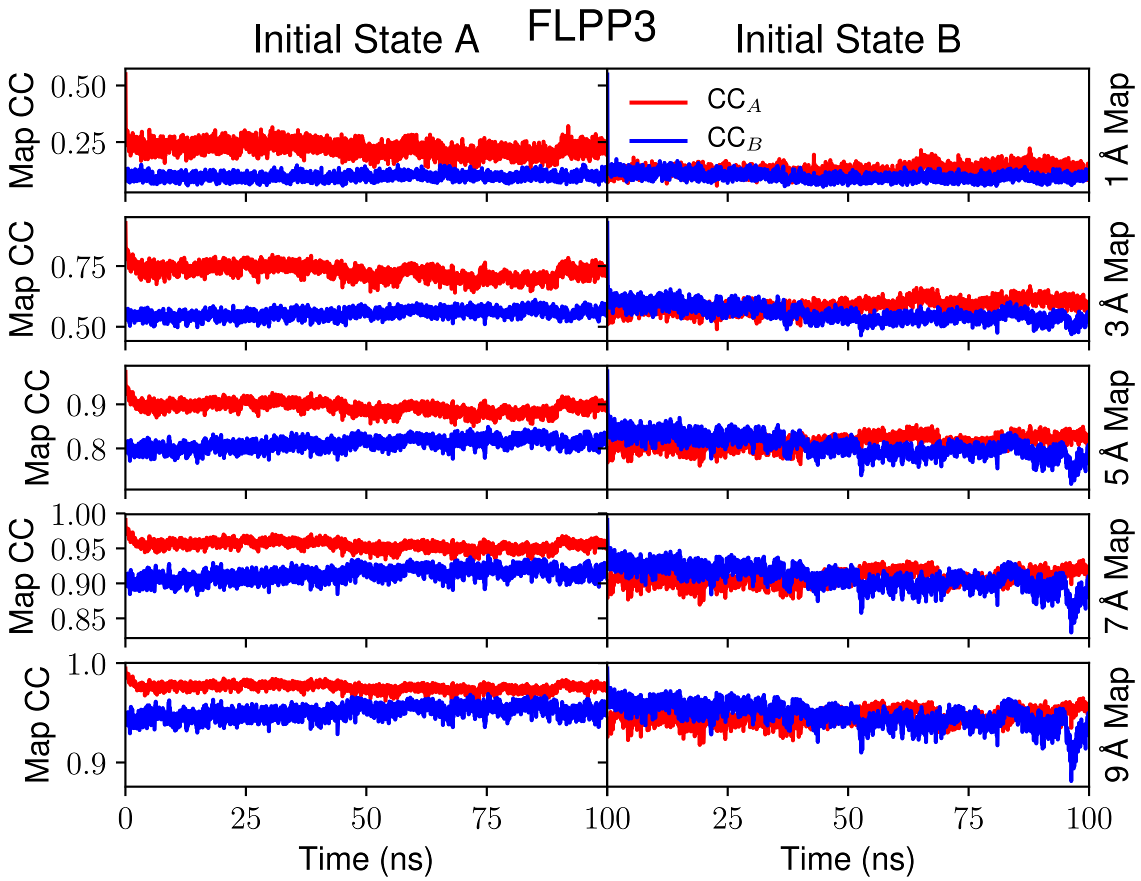

FIG. S6. Map cross correlation traces for equilibrium trajectories initialized from state A (left) or B (right) for FLPP3 with different map resolutions. $CC_A$ is the cross correlation with respect to the state A map. Likewise, $CC_B$ is the cross correlation with respect to the state B map.

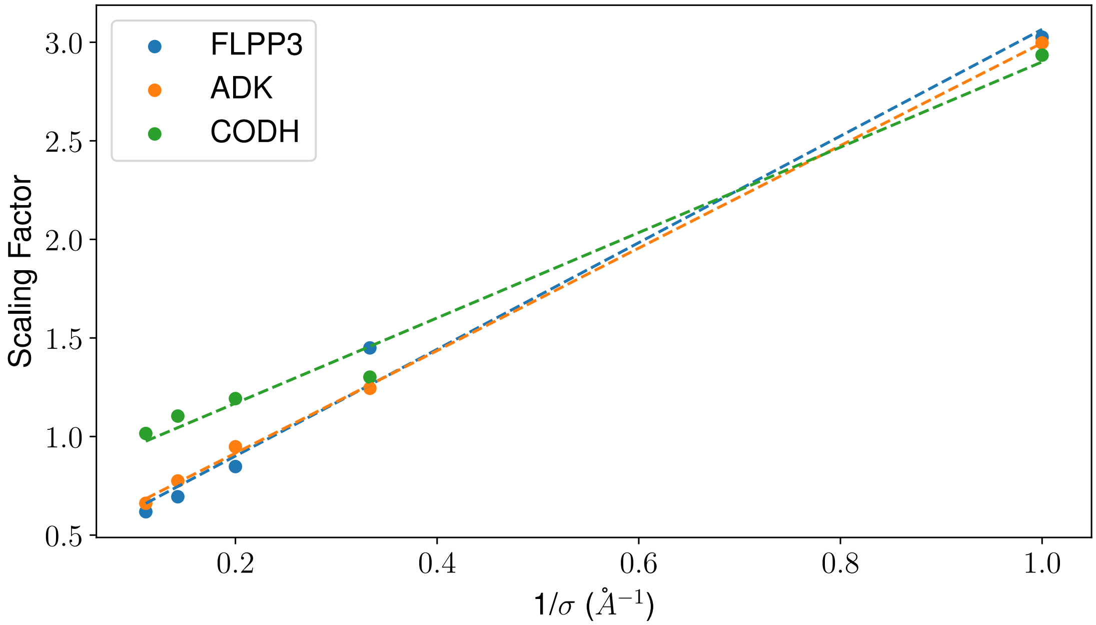

FIG. S7. Plot of the linear fits used for deriving $m$ and $b$ in Eq. S2 for each system under study. Where $\sigma$ is the resolution of the map and the scaling factor is the first factor presented in Eq.S2. The $m$ values were 2.71, 2.60 and 2.16 for FLPP3, ADK and CODH, respectively. The $b$ values were 0.36, 0.40 and 0.74 for FLPP3, ADK and CODH, respectively.

TABLE S2. Range for Multi-Map colvar in different size proteins at varying map resolutions.

| **Protein** | **Resolutions** | **Weight** [a] | **CC** [b] | **Actual** | **Predicted** [c] | **% error** |
|---|---|---|---|---|---|---|
| ADK (∼ 62,000 atom system) | 1 Å | 21883.327 | 0.081 | 6.03E+04 | 6.03E+04 | 0.03 |
| | 3 Å | 21883.327 | 0.414 | 1.60E+04 | 1.62E+04 | -1.42 |
| | 5 Å | 11577.884 | 0.619 | 4.18E+03 | 4.04E+03 | 3.42 |
| | 7 Å | 11577.884 | 0.713 | 2.58E+03 | 2.55E+03 | 0.97 |
| | 9 Å | 11577.884 | 0.763 | 1.82E+03 | 1.88E+03 | -3.50 |
| CODH (∼ 126,000 atom system) | 1 Å | 75811.850 | 0.124 | 1.95E+05 | 1.93E+05 | 1.18 |
| | 3 Å | 75811.850 | 0.507 | 4.87E+04 | 5.45E+04 | -11.99 |
| | 5 Å | 39348.019 | 0.722 | 1.30E+04 | 1.28E+04 | 1.99 |
| | 7 Å | 39348.019 | 0.829 | 7.44E+03 | 7.05E+03 | 5.32 |
| | 9 Å | 39348.019 | 0.881 | 4.77E+03 | 4.59E+03 | 3.85 |
| FLPP3 (∼ 42,000 atom system) | 1 Å | 11088.459 | 0.192 | 2.71E+04 | 2.75E+04 | -1.31 |
| | 3 Å | 11088.459 | 0.654 | 5.57E+03 | 4.85E+03 | 12.94 |
| | 5 Å | 5850.969 | 0.851 | 7.38E+02 | 7.85E+02 | -6.31 |
| | 7 Å | 5850.969 | 0.923 | 3.13E+02 | 3.36E+02 | -7.49 |
| | 9 Å | 5850.969 | 0.954 | 1.68E+02 | 1.79E+02 | -6.85 |

[a] Weight of protein atoms coupled to the density map.
[b] Correlation coefficient between model and density map.
[c] Prediction determined from Eq. S2

TABLE S3. Comparison of model quality using overall MolProbity[4] score between initial structure and model obtained from SMD, performed at different map resolutions.

| **Protein** | **A → B** | | | | | | **B → A** | | | | | |
|---|---|---|---|---|---|---|---|---|---|---|---|---|
| | **initial** | **1 Å** | **3 Å** | **5 Å** | **7 Å** | **9 Å** | **initial** | **1 Å** | **3 Å** | **5 Å** | **7 Å** | **9 Å** |
| ADK | 2.60 | 1.13 | 0.87 | 1.28 | 1.36 | 1.13 | 1.35 | 1.01 | 1.13 | 0.95 | 1.16 | 0.80 |
| CODH | 3.75 | 1.28 | 1.21 | 1.18 | 1.35 | 1.27 | 3.81 | 1.44 | 1.34 | 1.29 | 1.28 | 1.43 |
| FLPP3 | 0.71 | 1.69 | 1.63 | 0.72 | 1.18 | 1.26 | 1.82 | 1.40 | 1.35 | 1.36 | 1.05 | 1.03 |

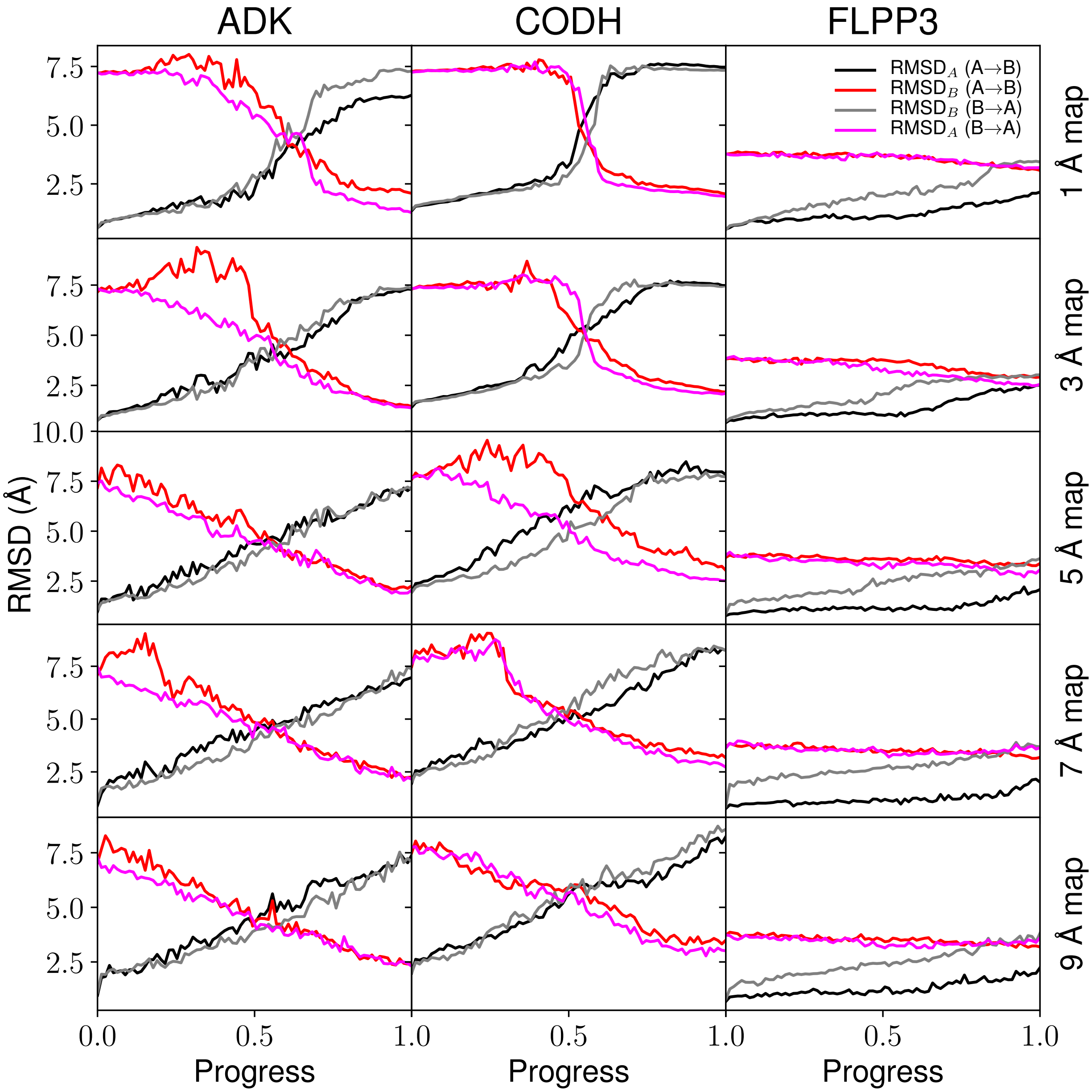

FIG. S8. Root mean square deviation values to either endpoint (RMSD$_A$ for state A, RMSD$_B$ for state B) for the two-state explicit solvent steered molecular dynamics trajectories.

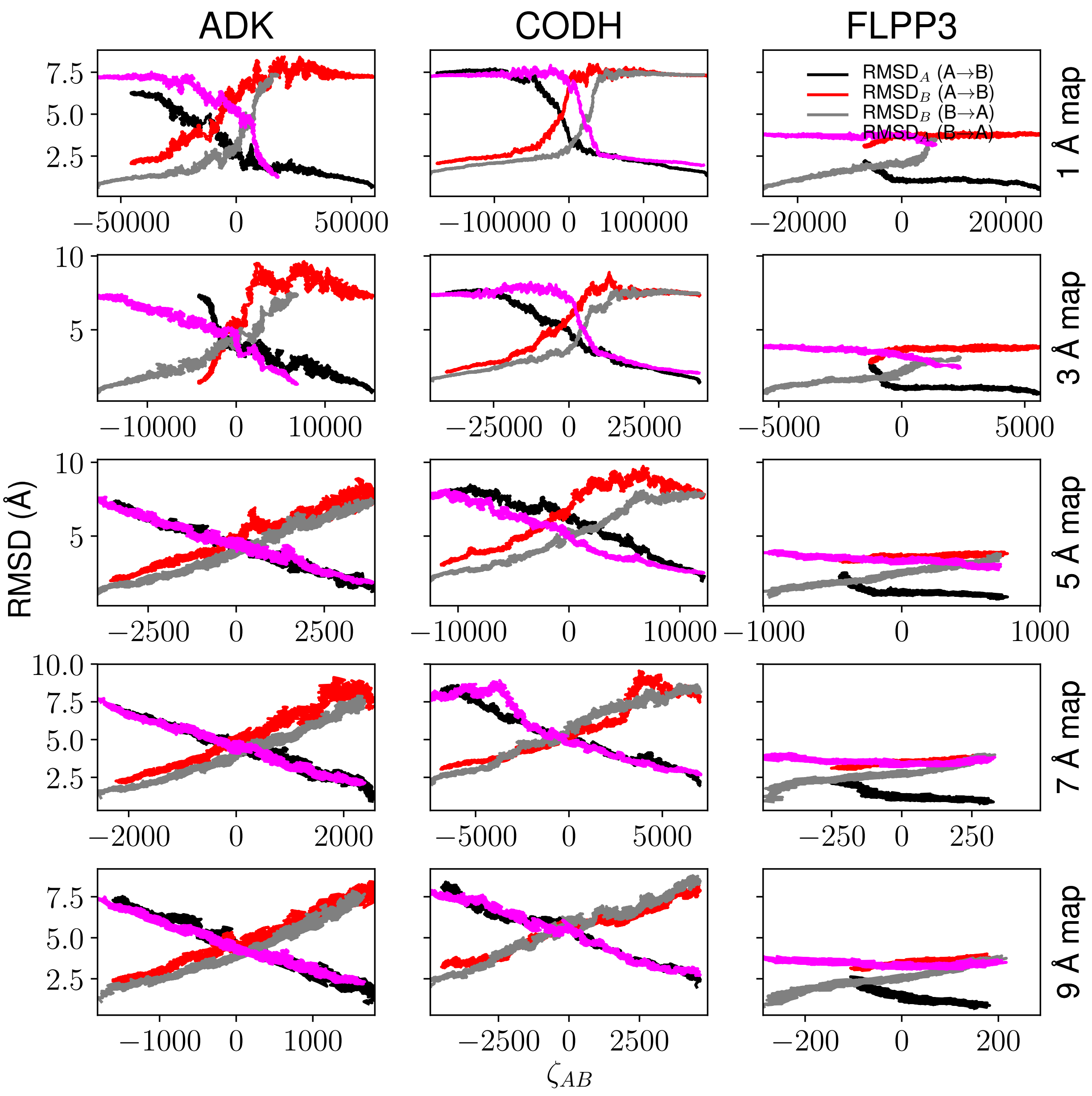

FIG. S9. Root mean square deviation values traced against the the $\zeta_{AB}$ values during the steered molecular dynamics trajectories. As in Fig. 4, each subplot simultaneously shows results for the A→B (black and red lines) and B→A (gray and fuchsia lines) transition. Colored lines highlight motion towards the target, as measured through the cross-correlation to the target density, while the black and gray lines represent the falling out of the density.

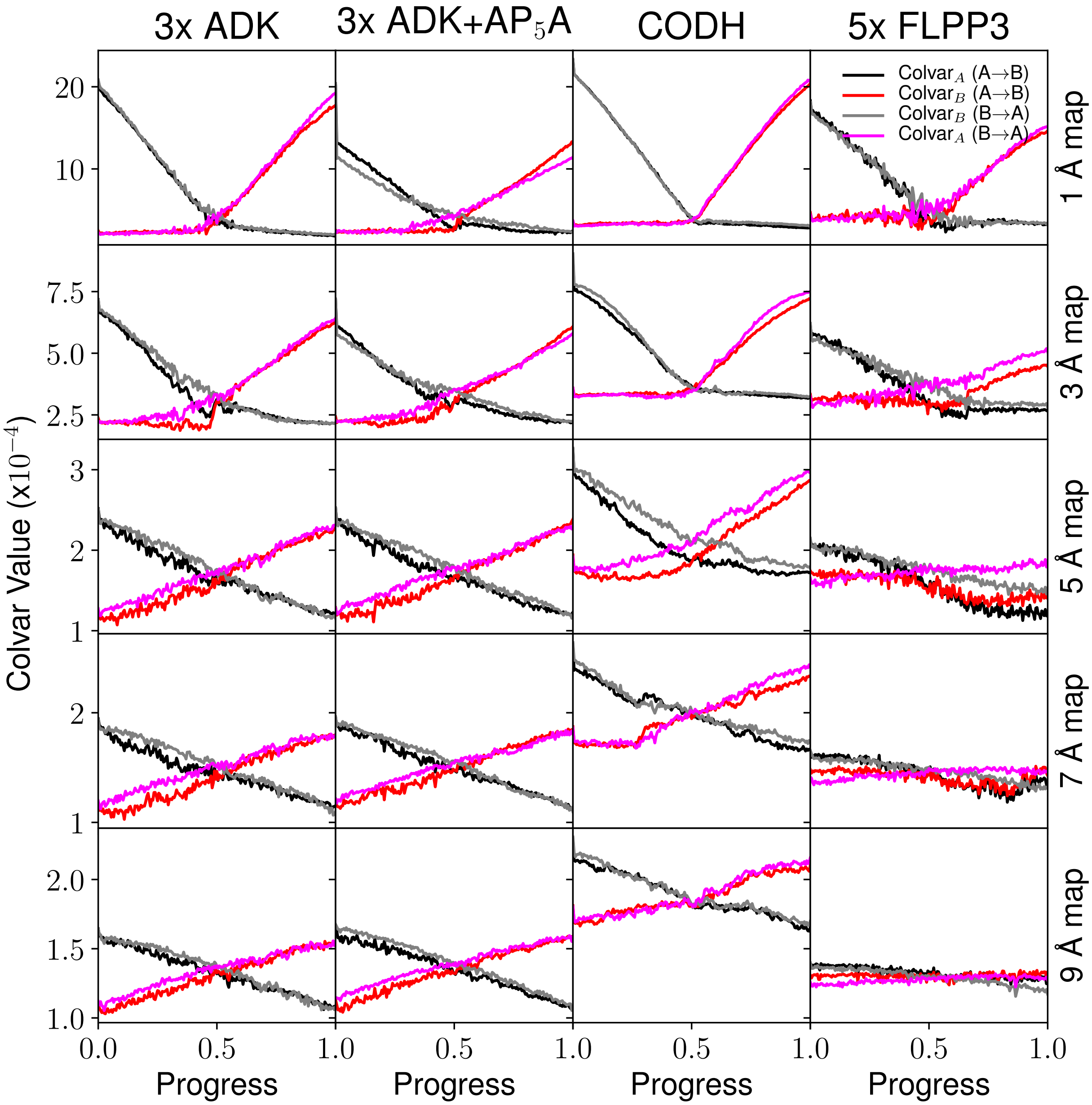

FIG. S10. Colvar values for correspondence to the initial and final maps (equivalent to $\zeta_A$ and $\zeta_B$ in Eq. 2) over different map resolutions during the explicit solvent steered molecular dynamics trajectories. Particularly for high resolution maps, the colvar value may fall quickly after time zero from its initial value.

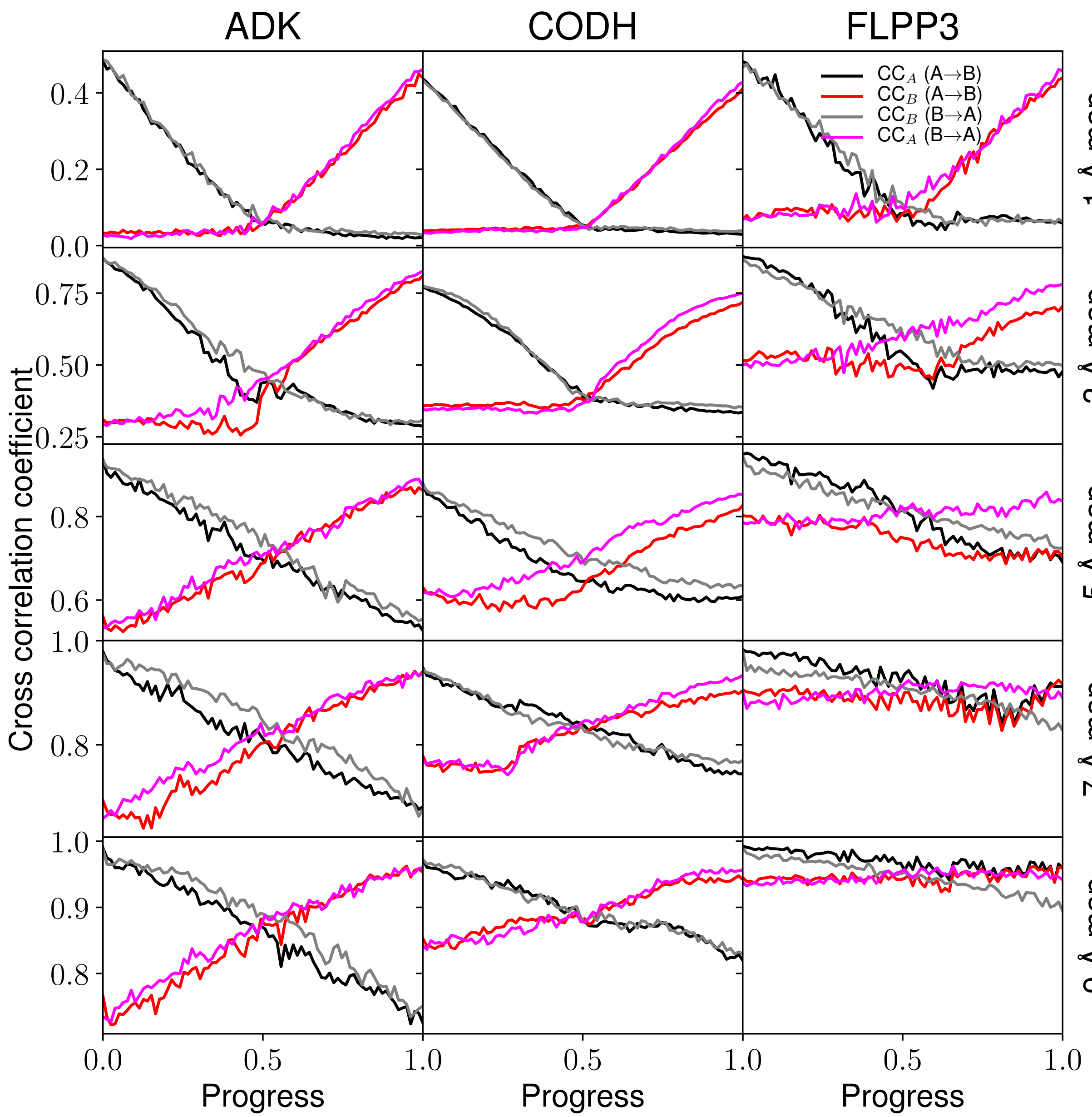

FIG. S11. Instantaneous map cross-correlation values to either endpoint (CC$_A$ for state A, CC$_B$ for state B) for the two-state explicit solvent steered molecular dynamics trajectories.

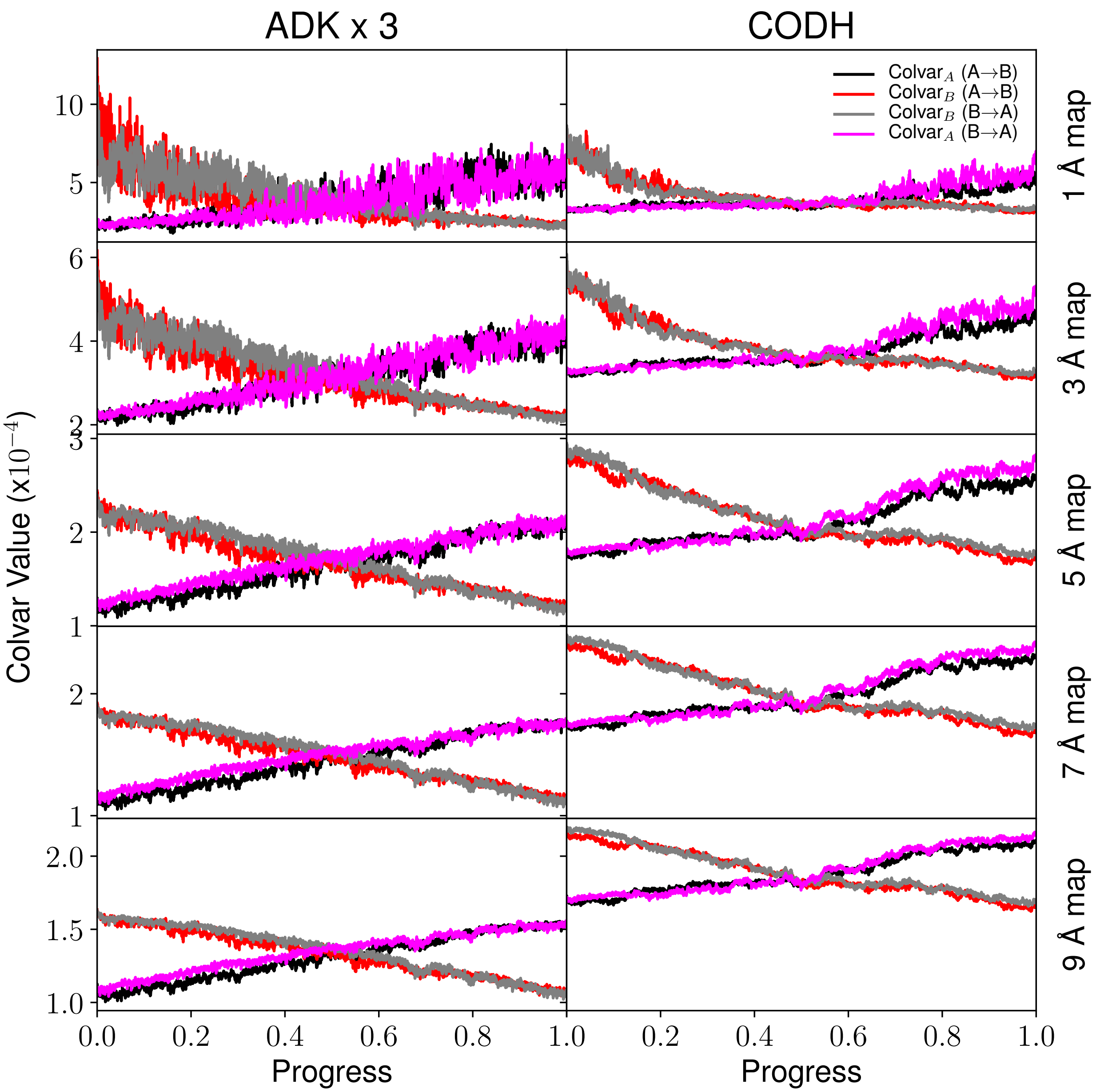

FIG. S12. Trajectories guided by the 9 Å map from state A to B and vice versa are measured with single map colvars $\zeta_A$ and $\zeta_B$ at resolutions 1, 3, 5, 7, and 9 Å.

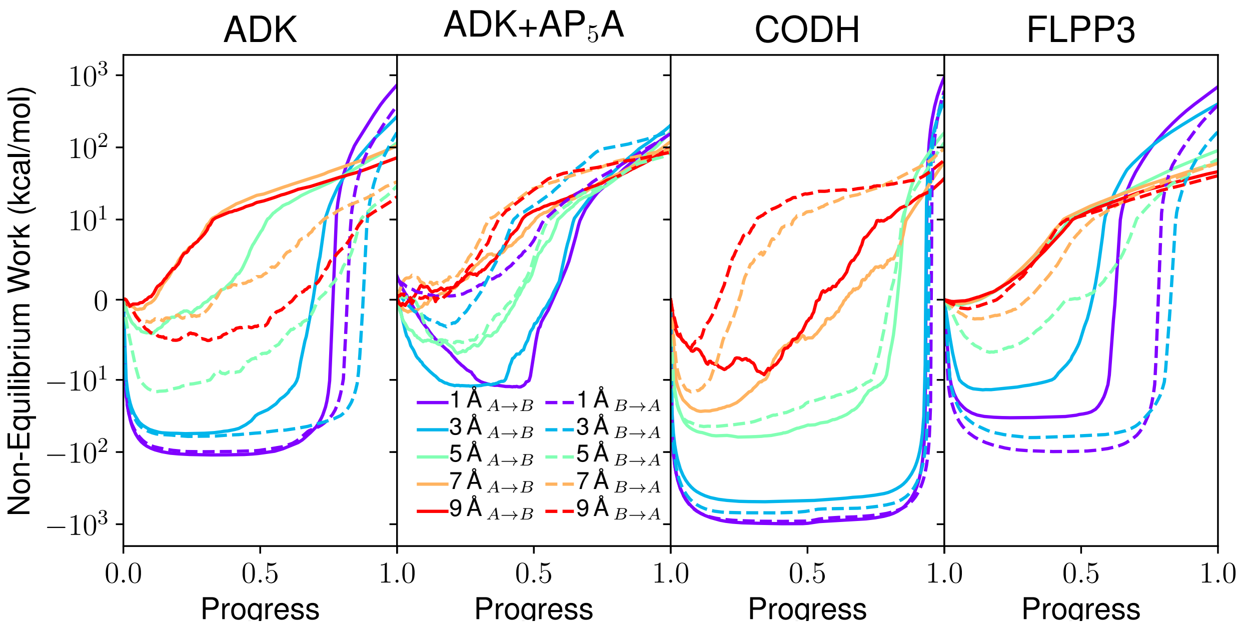

FIG. S13. Non-equilibrium work for the transitions in explicit solvent. Solid lines indicate the A→B transitions, while dashed lines indicate B→A transitions. Map resolution is indicated by line color, with bluer colors indicating high resolution maps, while redder colors indicate lower resolution maps. Note that due to the considerable variation in non-equilibrium work values, the plot is linear in the range (-10,10), and is plotted logarithmically outside of that range.

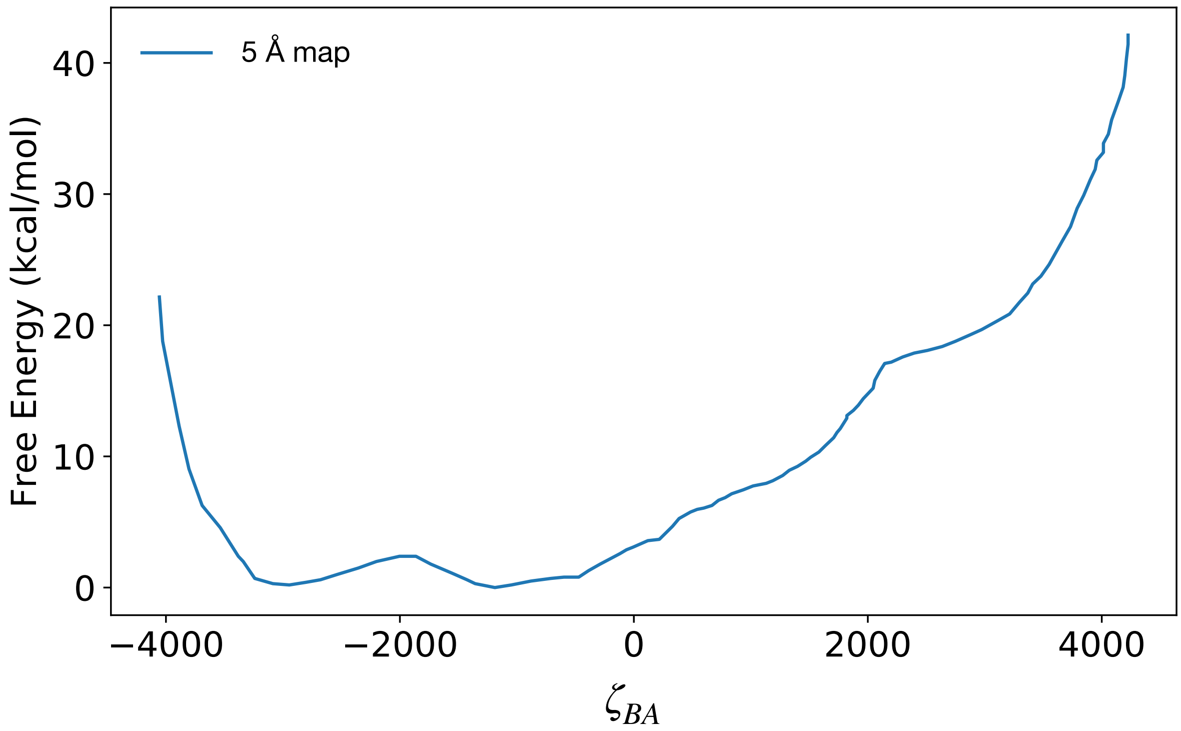

FIG. S14. Free energy profile for ADK in a vacuum using a 5 Å resolution maps for state A and B. At negative $\zeta_{BA}$ values the system is fit to state B and at positive $\zeta_{BA}$ values, the system is fit to state A. The PMF incorporates a total of 20 ns of BEUS sampling with 50 windows.

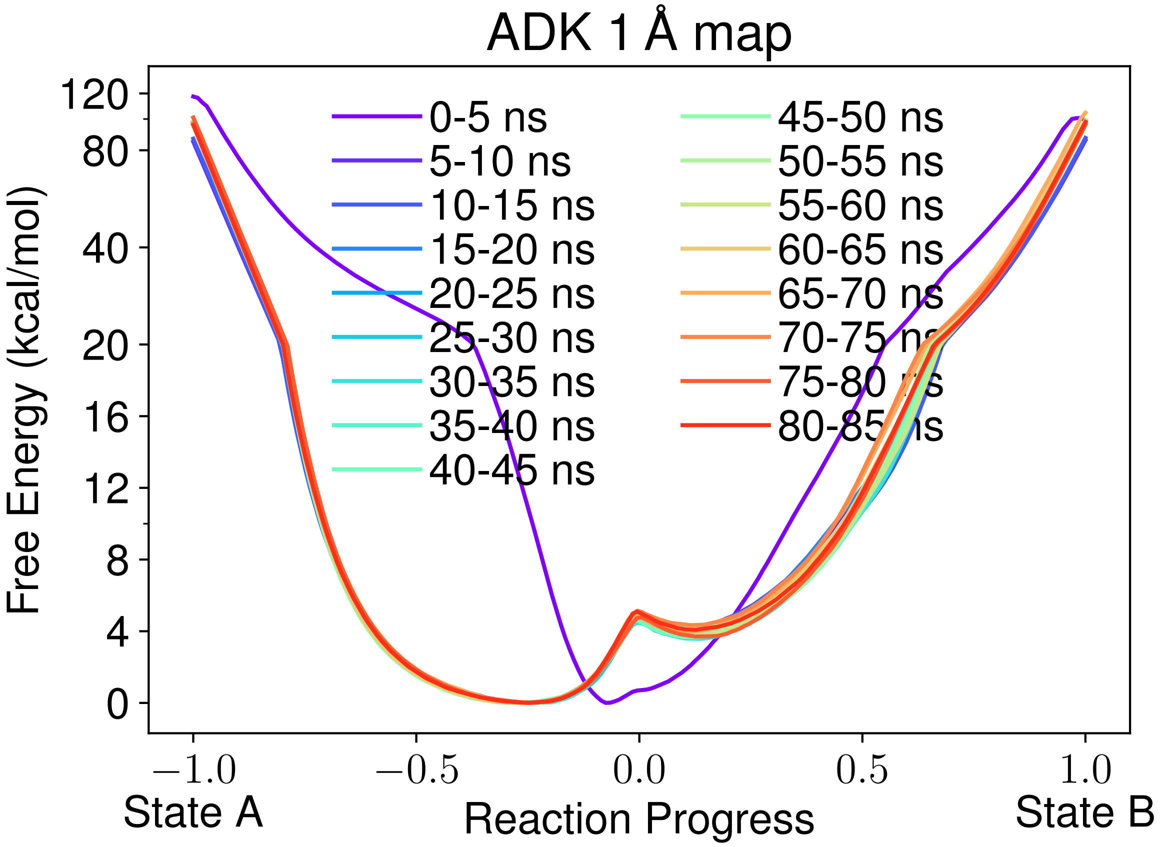

FIG. S15. Free energy convergence profiles for ADK using a 1 Å map resolution. Each profile is calculated from a 5 ns segment of the trajectory, to allow convergence and statistical deviation to be assessed. In some cases, the WHAM equations did not converge due to unsampled regions of space in the short simulation time. In these instances, those profiles are not reported. The x-axis has been scaled to reflect reaction progress, with values of -1 indicating a perfect map match with state A and +1 indicating map correspondence with state B.

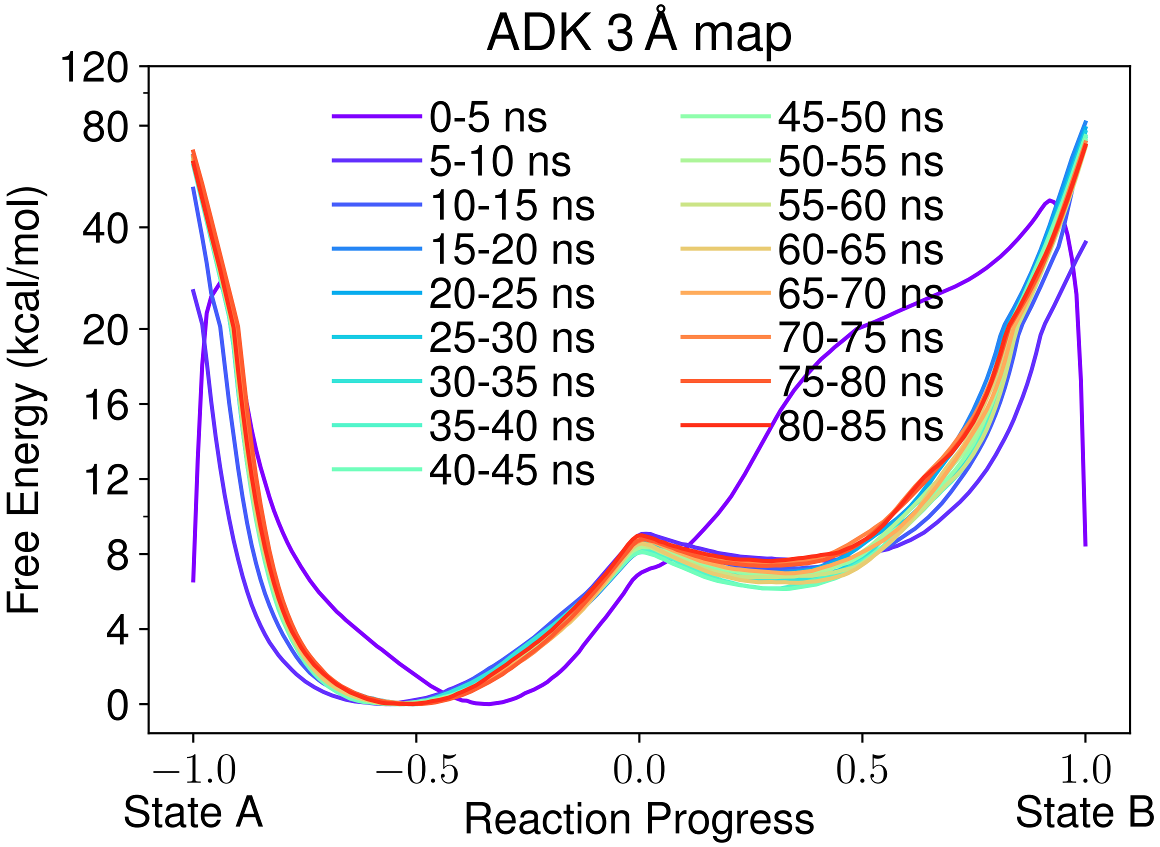

FIG. S16. Free energy convergence profiles for ADK using a 3 Å map resolution. Each profile is calculated from a 5 ns segment of the trajectory, to allow convergence and statistical deviation to be assessed. In some cases, the WHAM equations did not converge due to unsampled regions of space in the short simulation time. In these instances, those profiles are not reported. The x-axis has been scaled to reflect reaction progress, with values of -1 indicating a perfect map match with state A and +1 indicating map correspondence with state B.

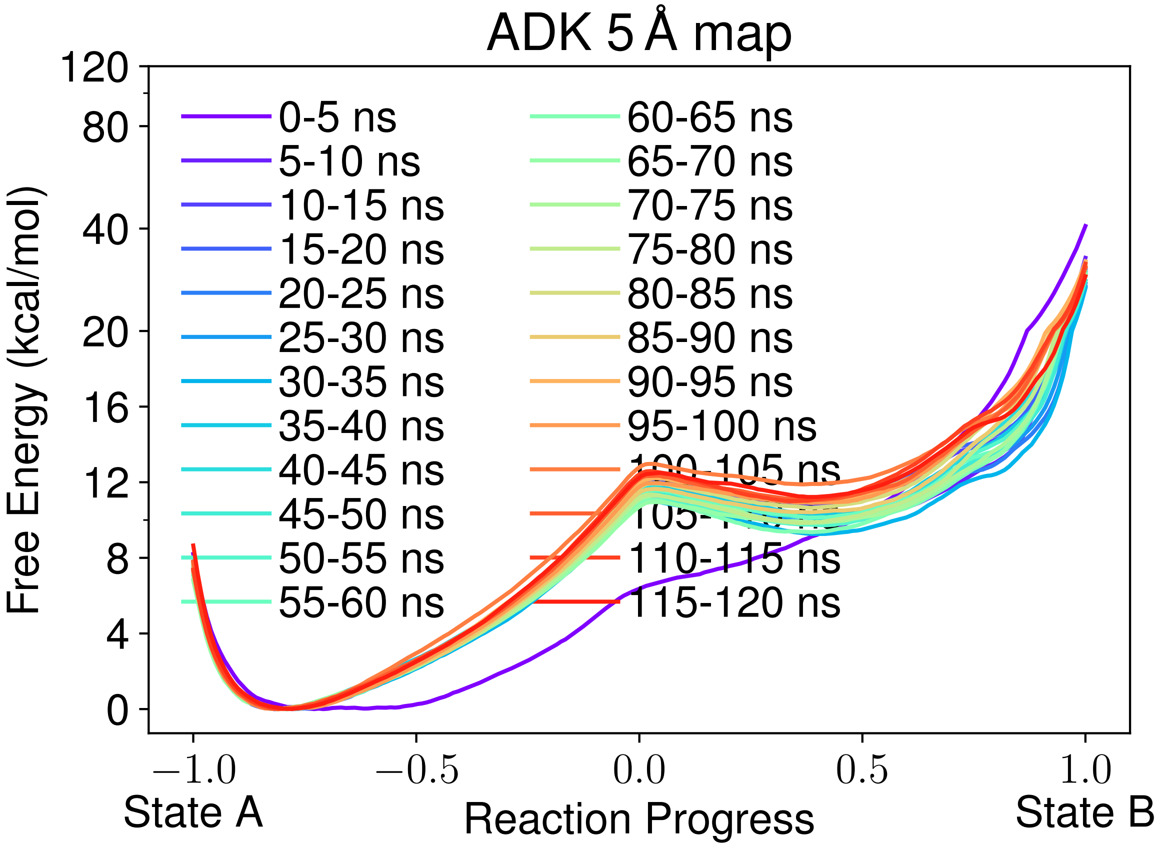

FIG. S17. Free energy convergence profiles for ADK using a 5 Å map resolution. Each profile is calculated from a 5 ns segment of the trajectory, to allow convergence and statistical deviation to be assessed. In some cases, the WHAM equations did not converge due to unsampled regions of space in the short simulation time. In these instances, those profiles are not reported. The x-axis has been scaled to reflect reaction progress, with values of -1 indicating a perfect map match with state A and +1 indicating map correspondence with state B.

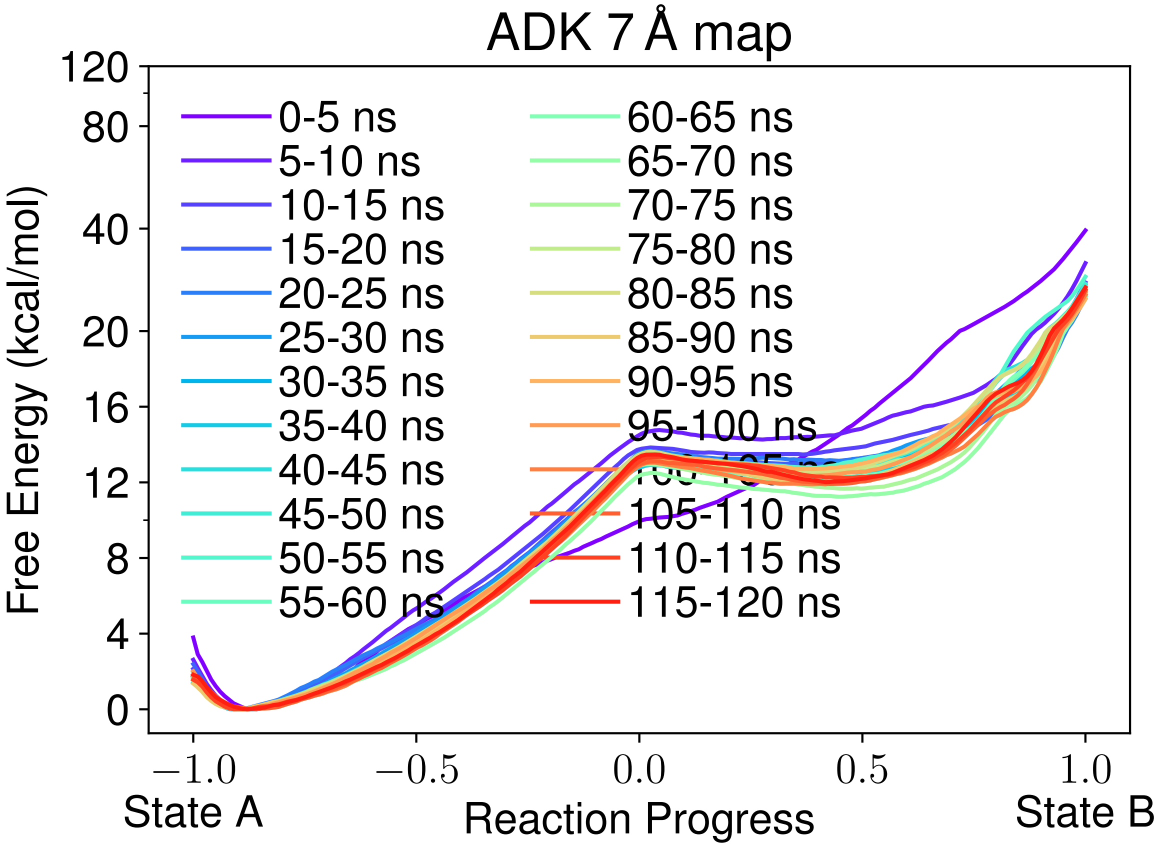

FIG. S18. Free energy convergence profiles for ADK using a 7 Å map resolution. Each profile is calculated from a 5 ns segment of the trajectory, to allow convergence and statistical deviation to be assessed. In some cases, the WHAM equations did not converge due to unsampled regions of space in the short simulation time. In these instances, those profiles are not reported. The x-axis has been scaled to reflect reaction progress, with values of -1 indicating a perfect map match with state A and +1 indicating map correspondence with state B.

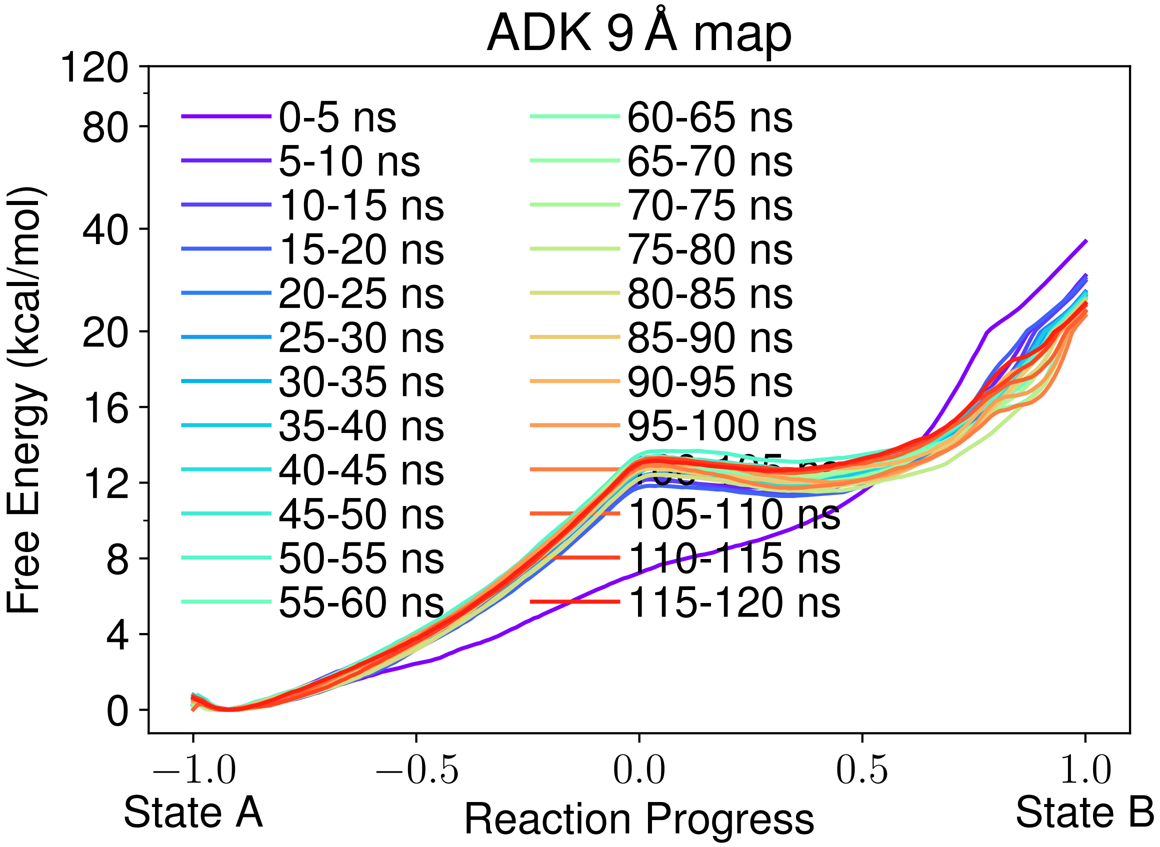

FIG. S19. Free energy convergence profiles for ADK using a 9 Å map resolution. Each profile is calculated from a 5 ns segment of the trajectory, to allow convergence and statistical deviation to be assessed. In some cases, the WHAM equations did not converge due to unsampled regions of space in the short simulation time. In these instances, those profiles are not reported. The x-axis has been scaled to reflect reaction progress, with values of -1 indicating a perfect map match with state A and +1 indicating map correspondence with state B.

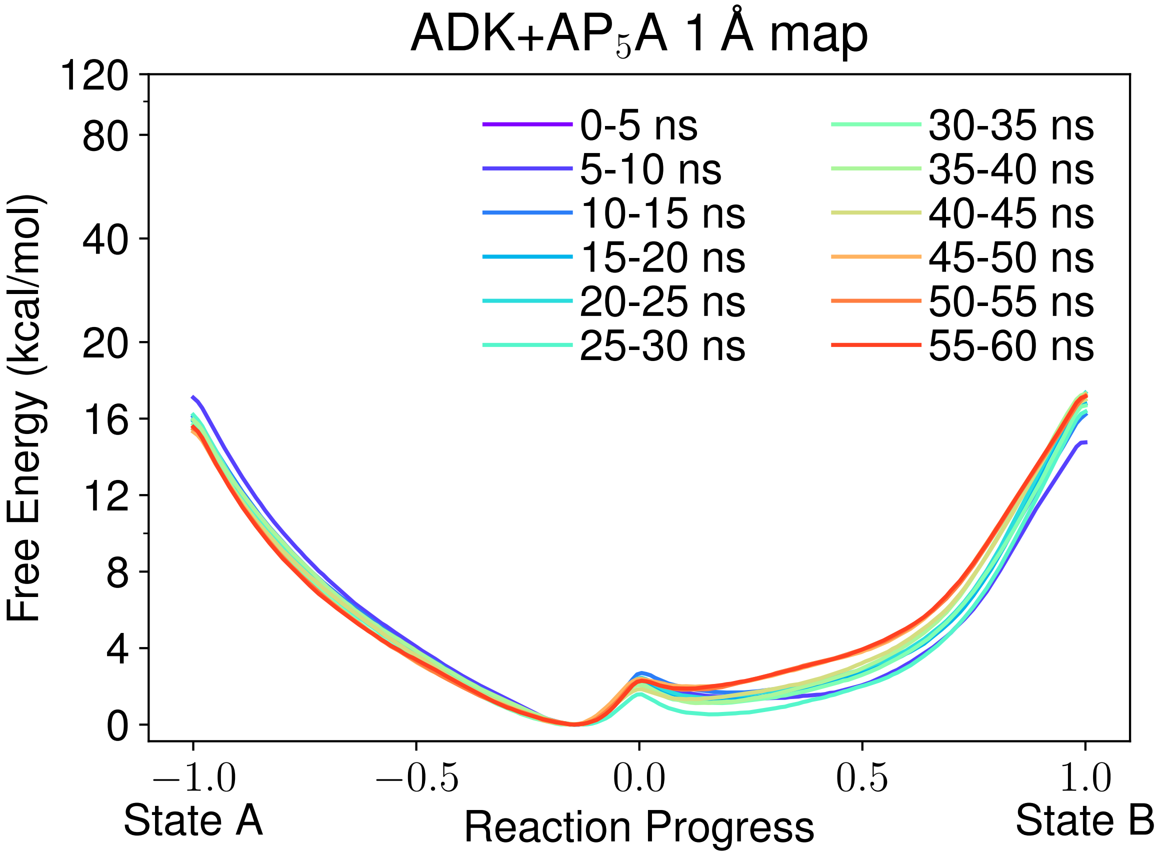

FIG. S20. Free energy convergence profiles for ADK-LIG using a 1 Å map resolution. Each profile is calculated from a 5 ns segment of the trajectory, to allow convergence and statistical deviation to be assessed. In some cases, the WHAM equations did not converge due to unsampled regions of space in the short simulation time. In these instances, those profiles are not reported. The x-axis has been scaled to reflect reaction progress, with values of -1 indicating a perfect map match with state A and +1 indicating map correspondence with state B.

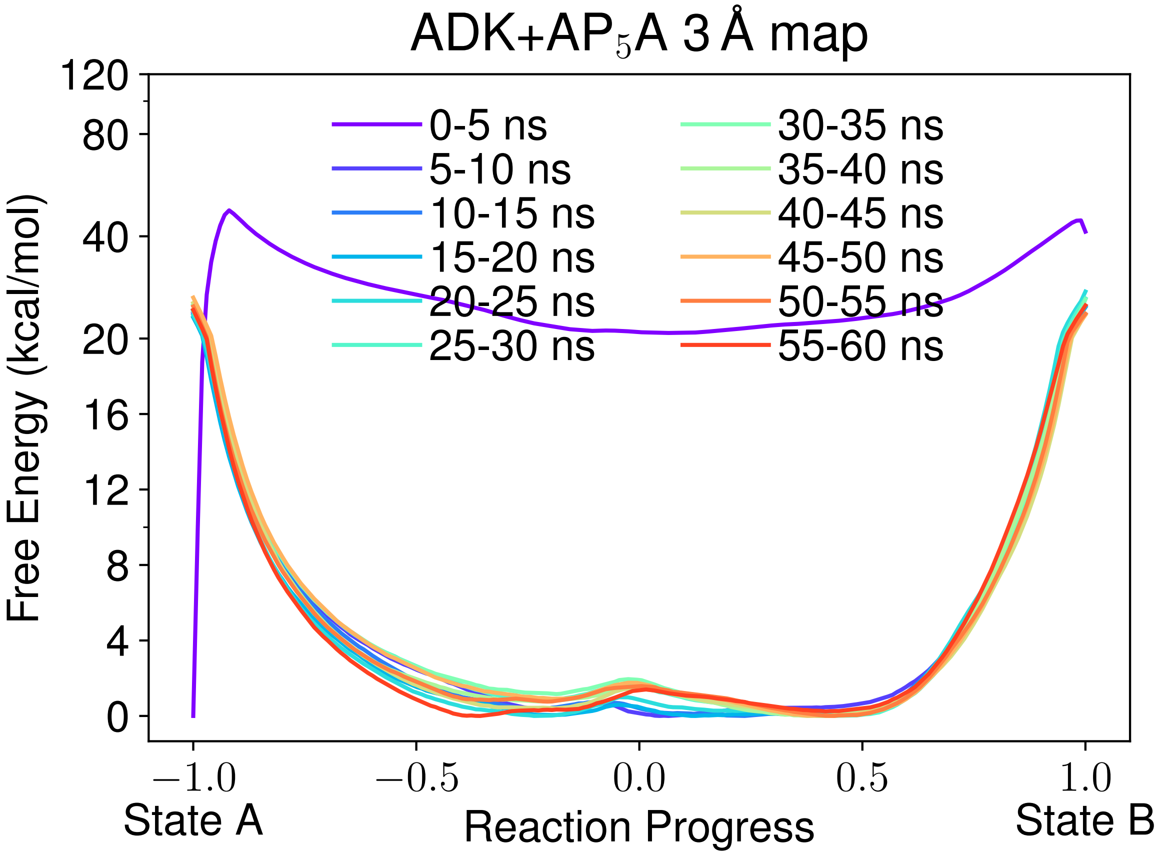

FIG. S21. Free energy convergence profiles for ADK-LIG using a 3 Å map resolution. Each profile is calculated from a 5 ns segment of the trajectory, to allow convergence and statistical deviation to be assessed. In some cases, the WHAM equations did not converge due to unsampled regions of space in the short simulation time. In these instances, those profiles are not reported. The x-axis has been scaled to reflect reaction progress, with values of -1 indicating a perfect map match with state A and +1 indicating map correspondence with state B.

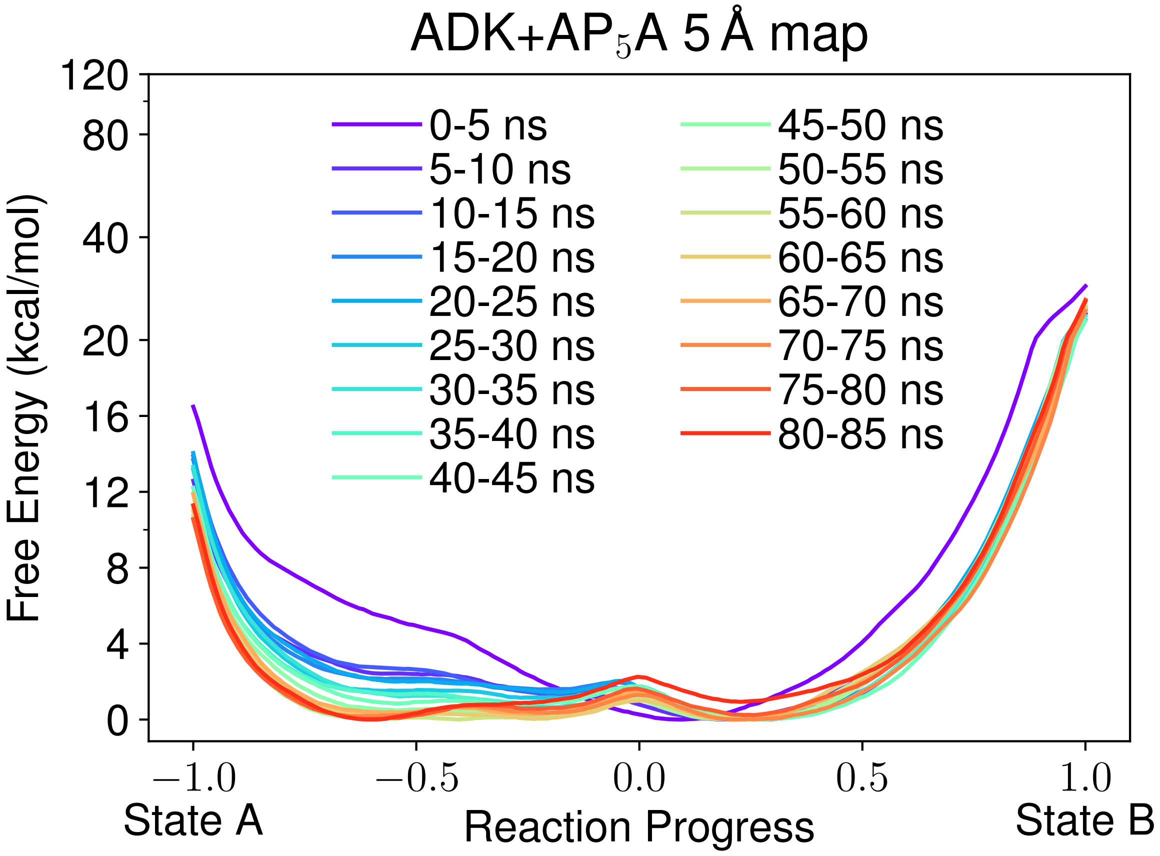

FIG. S22. Free energy convergence profiles for ADK-LIG using a 5 Å map resolution. Each profile is calculated from a 5 ns segment of the trajectory, to allow convergence and statistical deviation to be assessed. In some cases, the WHAM equations did not converge due to unsampled regions of space in the short simulation time. In these instances, those profiles are not reported. The x-axis has been scaled to reflect reaction progress, with values of -1 indicating a perfect map match with state A and +1 indicating map correspondence with state B.

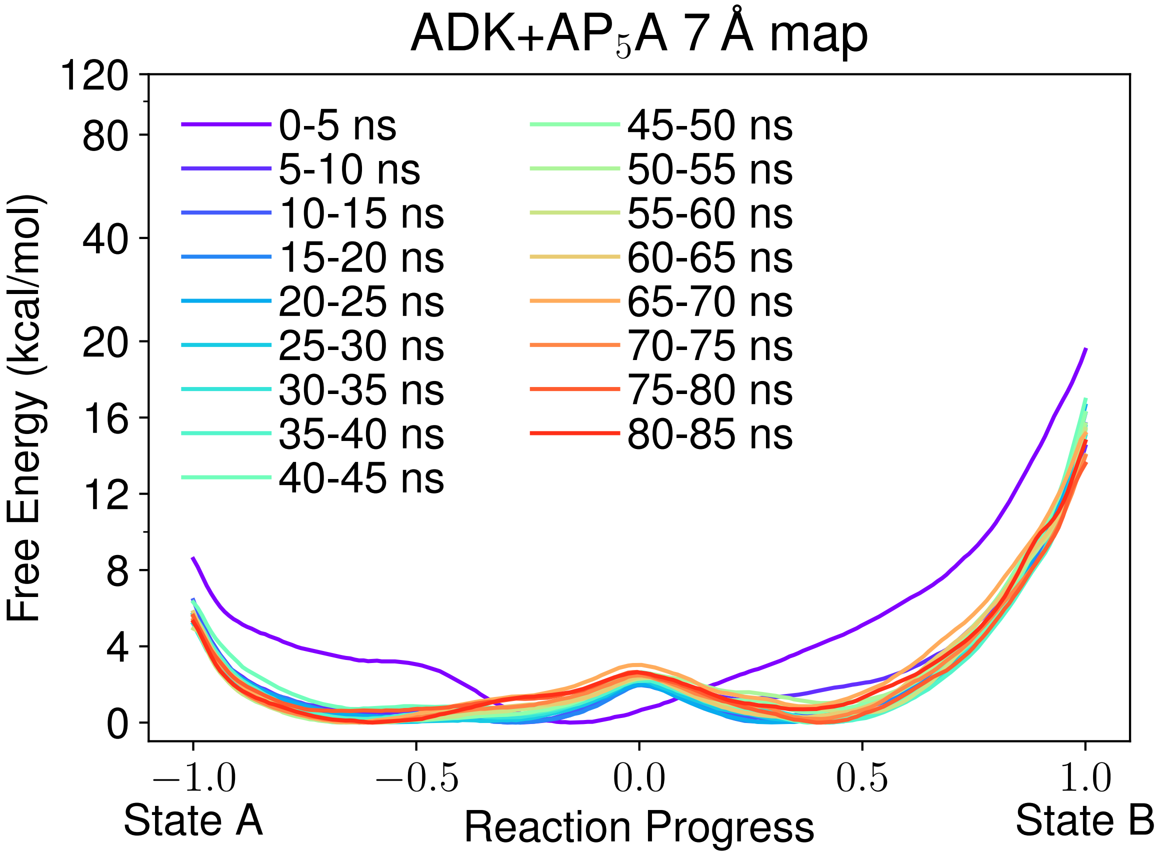

FIG. S23. Free energy convergence profiles for ADK-LIG using a 7 Å map resolution. Each profile is calculated from a 5 ns segment of the trajectory, to allow convergence and statistical deviation to be assessed. In some cases, the WHAM equations did not converge due to unsampled regions of space in the short simulation time. In these instances, those profiles are not reported. The x-axis has been scaled to reflect reaction progress, with values of -1 indicating a perfect map match with state A and +1 indicating map correspondence with state B.

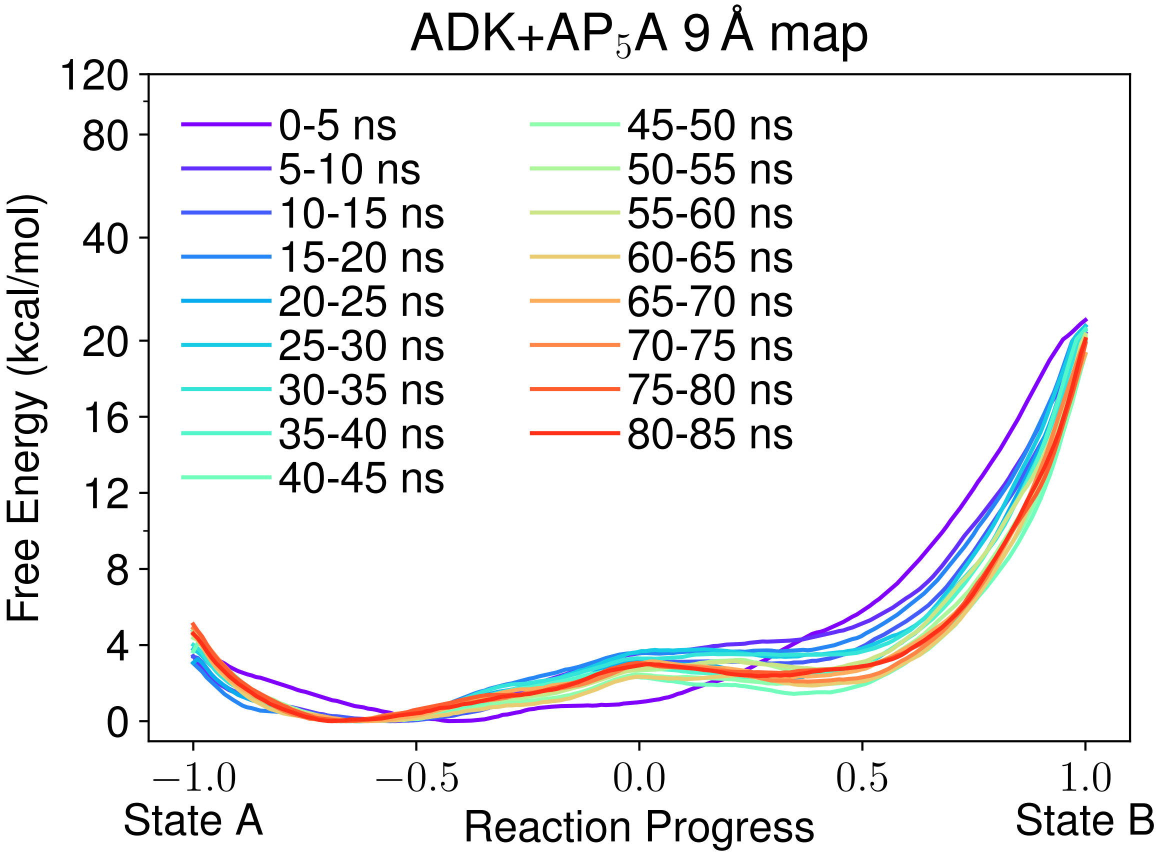

FIG. S24. Free energy convergence profiles for ADK-LIG using a 9 Å map resolution. Each profile is calculated from a 5 ns segment of the trajectory, to allow convergence and statistical deviation to be assessed. In some cases, the WHAM equations did not converge due to unsampled regions of space in the short simulation time. In these instances, those profiles are not reported. The x-axis has been scaled to reflect reaction progress, with values of -1 indicating a perfect map match with state A and +1 indicating map correspondence with state B.

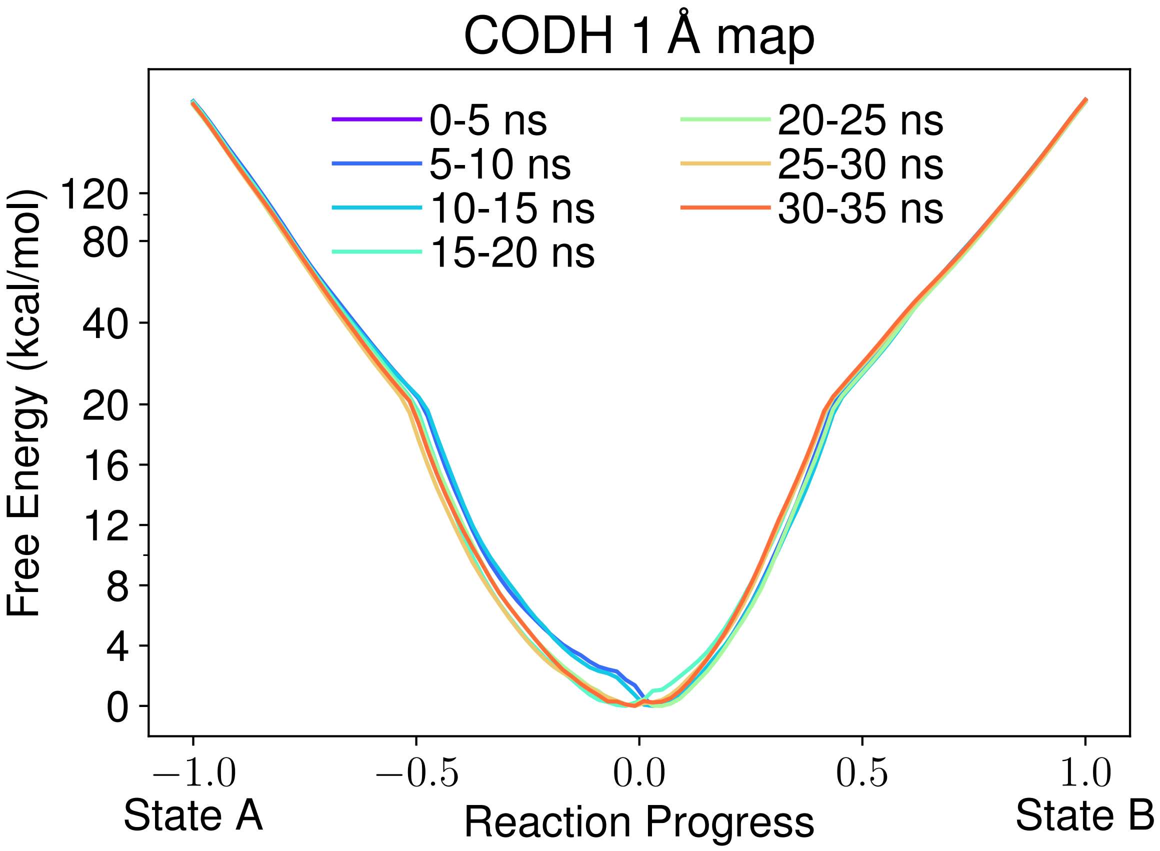

FIG. S25. Free energy convergence profiles for CODH using a 1 Å map resolution. Each profile is calculated from a 5 ns segment of the trajectory, to allow convergence and statistical deviation to be assessed. In some cases, the WHAM equations did not converge due to unsampled regions of space in the short simulation time. In these instances, those profiles are not reported. The x-axis has been scaled to reflect reaction progress, with values of -1 indicating a perfect map match with state A and +1 indicating map correspondence with state B.

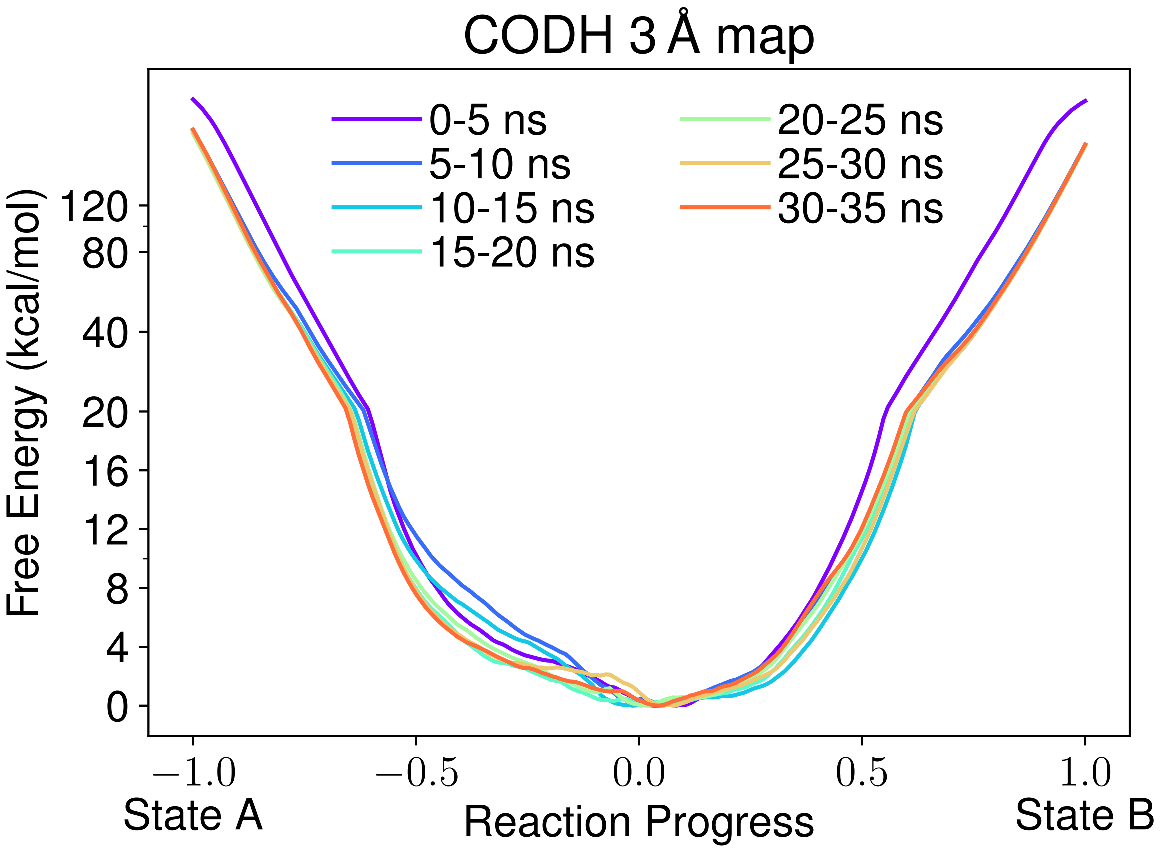

FIG. S26. Free energy convergence profiles for CODH using a 3 Å map resolution. Each profile is calculated from a 5 ns segment of the trajectory, to allow convergence and statistical deviation to be assessed. In some cases, the WHAM equations did not converge due to unsampled regions of space in the short simulation time. In these instances, those profiles are not reported. The x-axis has been scaled to reflect reaction progress, with values of -1 indicating a perfect map match with state A and +1 indicating map correspondence with state B.

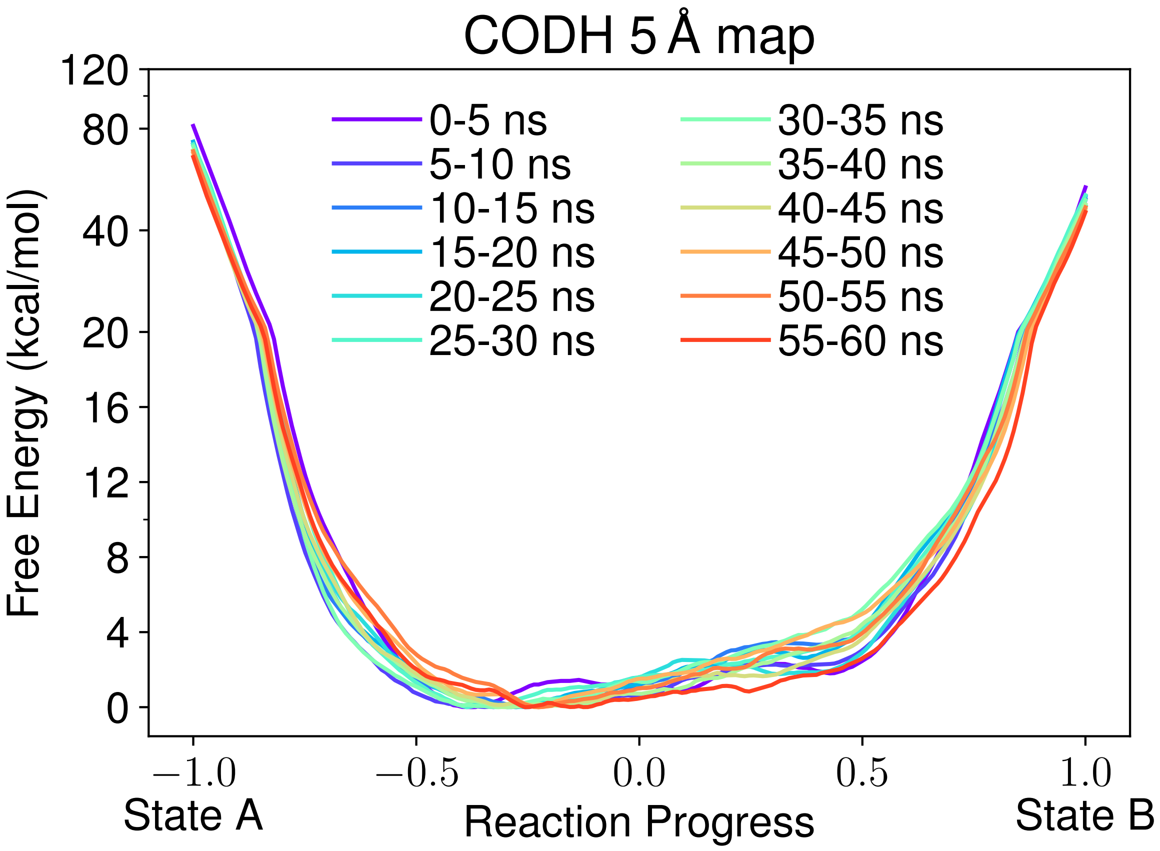

FIG. S27. Free energy convergence profiles for CODH using a 5 Å map resolution. Each profile is calculated from a 5 ns segment of the trajectory, to allow convergence and statistical deviation to be assessed. In some cases, the WHAM equations did not converge due to unsampled regions of space in the short simulation time. In these instances, those profiles are not reported. The x-axis has been scaled to reflect reaction progress, with values of -1 indicating a perfect map match with state A and +1 indicating map correspondence with state B.

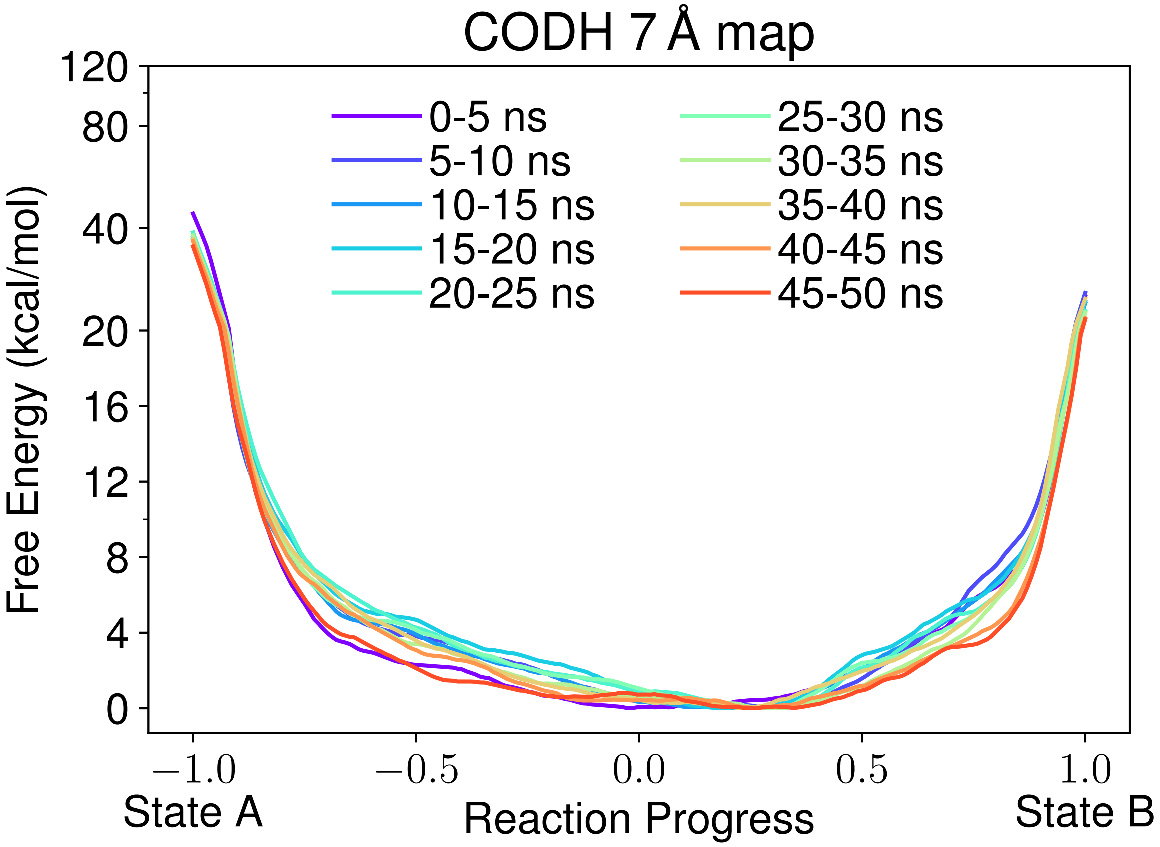

FIG. S28. Free energy convergence profiles for CODH using a 7 Å map resolution. Each profile is calculated from a 5 ns segment of the trajectory, to allow convergence and statistical deviation to be assessed. In some cases, the WHAM equations did not converge due to unsampled regions of space in the short simulation time. In these instances, those profiles are not reported. The x-axis has been scaled to reflect reaction progress, with values of -1 indicating a perfect map match with state A and +1 indicating map correspondence with state B.

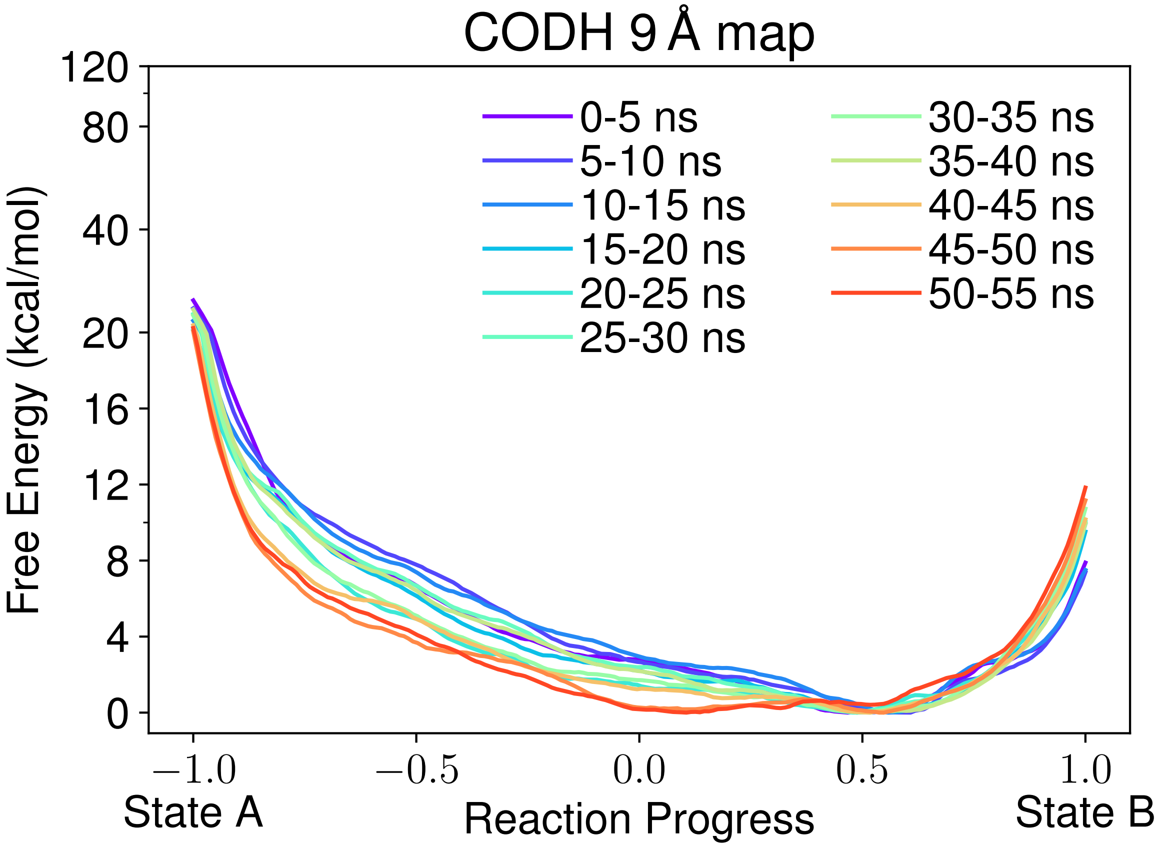

FIG. S29. Free energy convergence profiles for CODH using a 9 Å map resolution. Each profile is calculated from a 5 ns segment of the trajectory, to allow convergence and statistical deviation to be assessed. In some cases, the WHAM equations did not converge due to unsampled regions of space in the short simulation time. In these instances, those profiles are not reported. The x-axis has been scaled to reflect reaction progress, with values of -1 indicating a perfect map match with state A and +1 indicating map correspondence with state B.

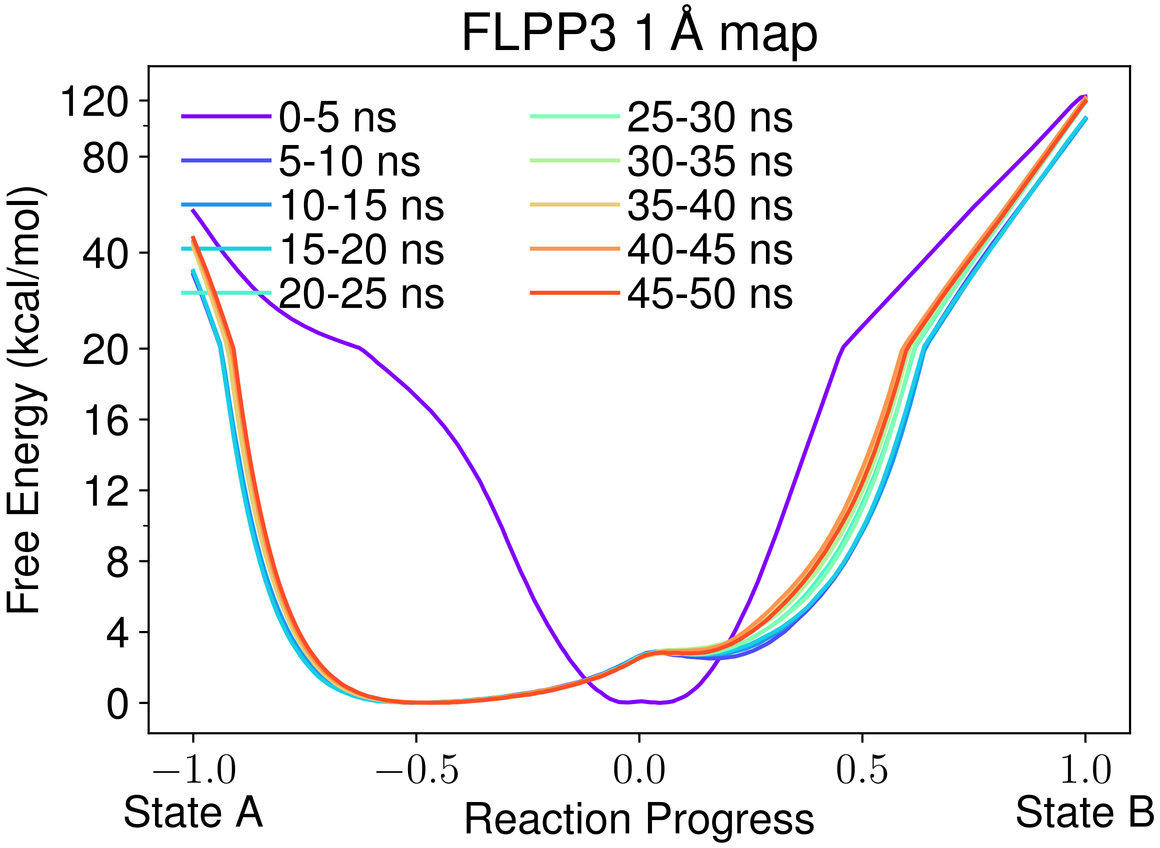

FIG. S30. Free energy convergence profiles for FLPP3 using a 1 Å map resolution. Each profile is calculated from a 5 ns segment of the trajectory, to allow convergence and statistical deviation to be assessed. In some cases, the WHAM equations did not converge due to unsampled regions of space in the short simulation time. In these instances, those profiles are not reported. The x-axis has been scaled to reflect reaction progress, with values of -1 indicating a perfect map match with state A and +1 indicating map correspondence with state B.

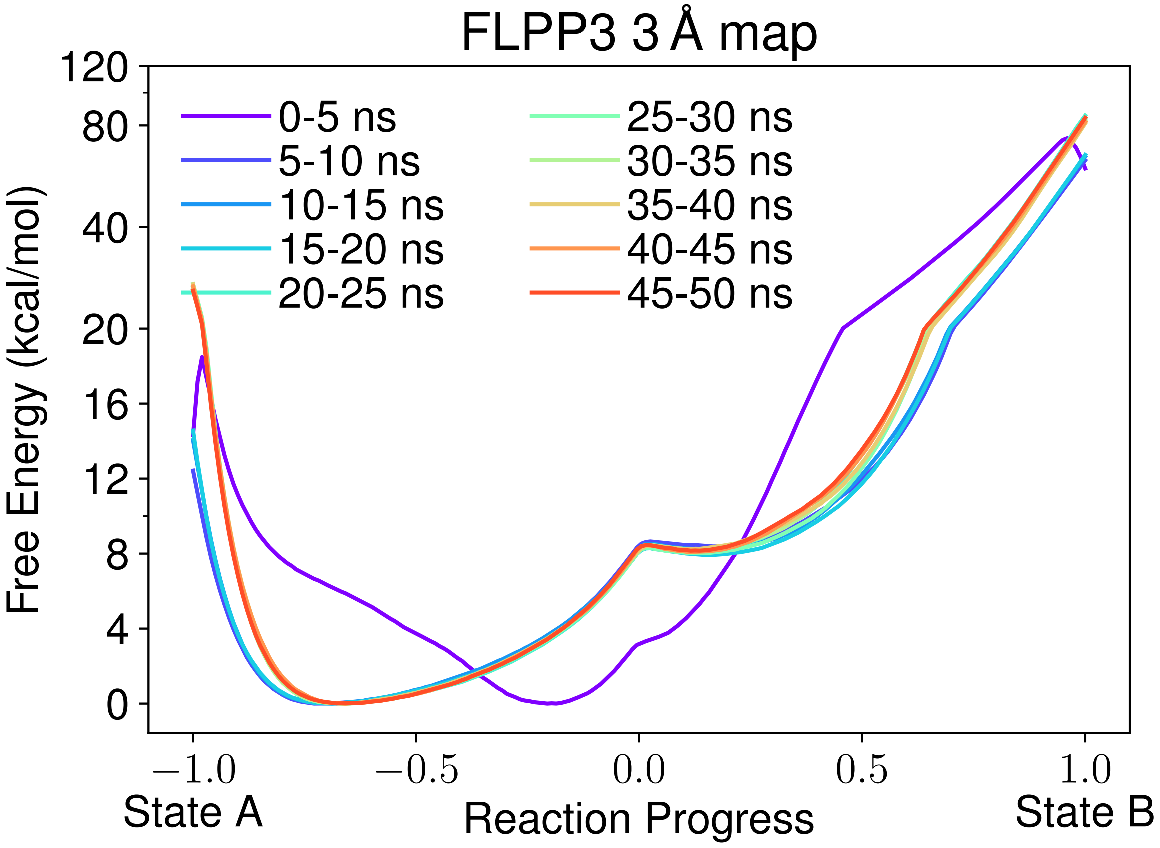

FIG. S31. Free energy convergence profiles for FLPP3 using a 3 Å map resolution. Each profile is calculated from a 5 ns segment of the trajectory, to allow convergence and statistical deviation to be assessed. In some cases, the WHAM equations did not converge due to unsampled regions of space in the short simulation time. In these instances, those profiles are not reported. The x-axis has been scaled to reflect reaction progress, with values of -1 indicating a perfect map match with state A and +1 indicating map correspondence with state B.

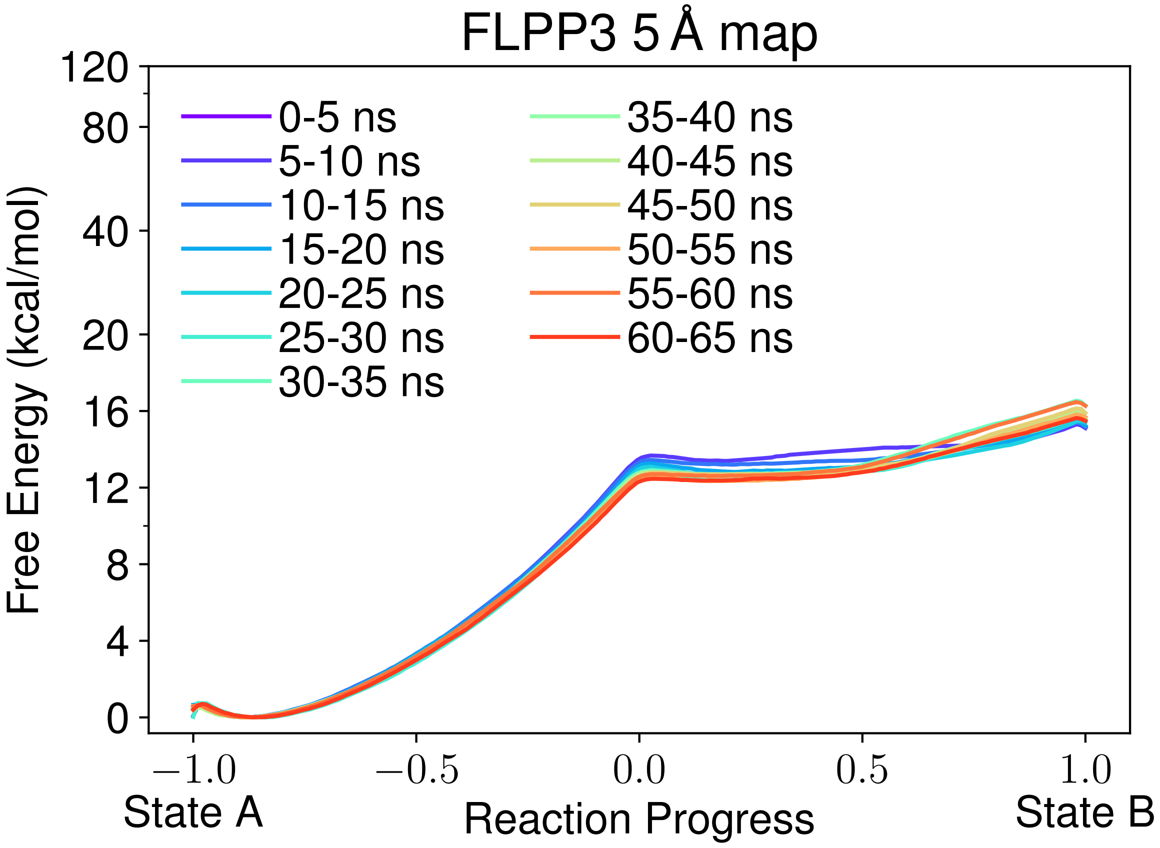

FIG. S32. Free energy convergence profiles for FLPP3 using a 5 Å map resolution. Each profile is calculated from a 5 ns segment of the trajectory, to allow convergence and statistical deviation to be assessed. In some cases, the WHAM equations did not converge due to unsampled regions of space in the short simulation time. In these instances, those profiles are not reported. The x-axis has been scaled to reflect reaction progress, with values of -1 indicating a perfect map match with state A and +1 indicating map correspondence with state B.

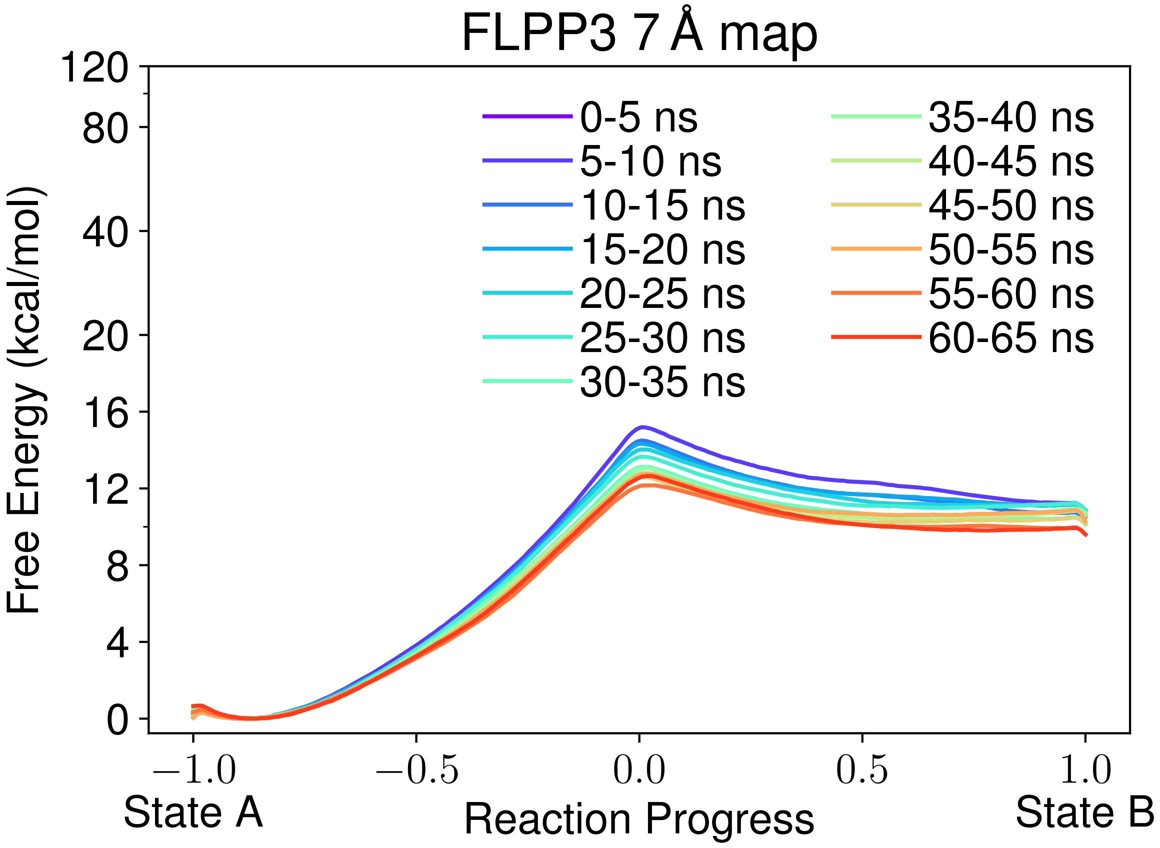

FIG. S33. Free energy convergence profiles for FLPP3 using a 7 Å map resolution. Each profile is calculated from a 5 ns segment of the trajectory, to allow convergence and statistical deviation to be assessed. In some cases, the WHAM equations did not converge due to unsampled regions of space in the short simulation time. In these instances, those profiles are not reported. The x-axis has been scaled to reflect reaction progress, with values of -1 indicating a perfect map match with state A and +1 indicating map correspondence with state B.

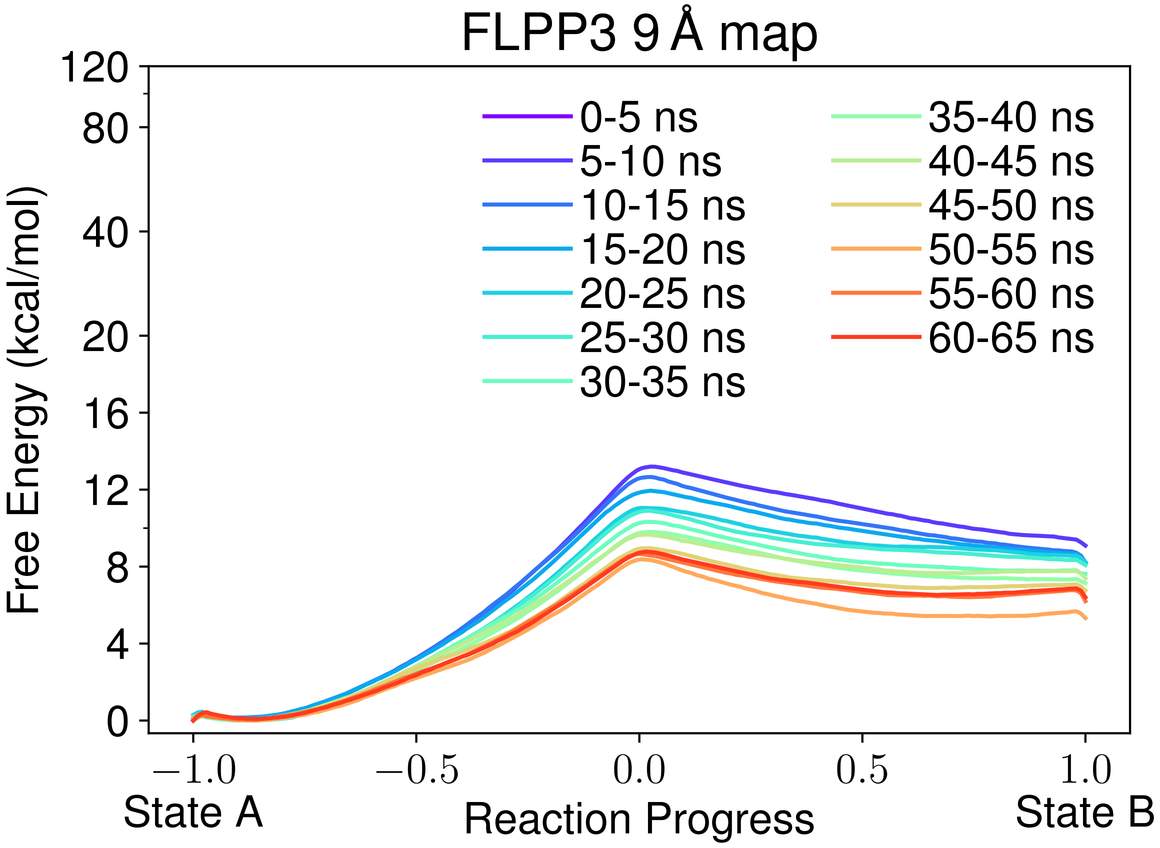

FIG. S34. Free energy convergence profiles for FLPP3 using a 9 Å map resolution. Each profile is calculated from a 5 ns segment of the trajectory, to allow convergence and statistical deviation to be assessed. In some cases, the WHAM equations did not converge due to unsampled regions of space in the short simulation time. In these instances, those profiles are not reported. The x-axis has been scaled to reflect reaction progress, with values of -1 indicating a perfect map match with state A and +1 indicating map correspondence with state B.
